# Expertise modifies neural responses to causal inference during language comprehension

**DOI:** 10.64898/2026.11.01.763546

**Authors:** Miriam Hauptman, Marina Bedny

**Affiliations:** Department of Psychological & Brain Sciences, Johns Hopkins University, Baltimore, MD, USA

**Keywords:** language, causal knowledge, fMRI, expertise, semantics

## Abstract

Linguistic communication relies on shared common ground, yet different individuals in a community have specialized semantic knowledge. As a key example, physicians acquire rich knowledge about the causes of illness. An open question is whether such “causal expertise” influences the neural basis of language comprehension. Here we tested the hypothesis that when understanding everyday language like, “Holly used a grimy faucet during her overseas plane ride. Now she has the stomach flu,” medical experts retrieve richer representations in semantic networks of the brain. Physicians (n=16) and age- and education-matched controls (n=16) undergoing fMRI read two-sentence vignettes that elicited implicit causal inferences about illness or object mechanics as well as control vignettes that did not elicit causal inferences.

Vignettes were linguistically matched across conditions and were accessible to experts and non-experts alike. Across both groups, causal vignettes about illness engaged a temporoparietal semantic network previously implicated in thinking about animates, i.e., people and animals (precuneus and left temporoparietal junction), whereas causal vignettes about object mechanics engaged a semantic network associated with places and objects. Critically, physicians’ responses in the animacy network were more robust and self-similar, suggesting the activation of richer semantic knowledge. Medical expertise effects were specific to biological inferences rather than mechanical or mental state inferences. By contrast, individually localized frontotemporal language and frontoparietal reasoning networks did not respond preferentially to causally connected sentences or show effects of expertise. Our findings suggest that causal expertise alters neural responses to causal inferences that unfold automatically during language comprehension.

**Significance statement:** Shared knowledge is necessary for communication, but what we know about the world varies significantly across individuals. Medical expertise offers a classic example. Relative to other people, medical experts develop a richer understanding of what causes illness. Here we asked whether becoming a “causal expert” alters neural responses during language comprehension. We find that physicians show enhanced responses to causal inferences about illness elicited by language. This expertise effect is specific to semantic networks of the brain that represent knowledge about animate entities, such as people and animals. Our findings suggest that content-specific semantic networks enable causal inferences during language comprehension and are modified by causal expertise, offering new insight into how conceptual variation shapes the neural basis of language.

## Introduction

Human societies are characterized by a division of intellectual labor: individuals acquire specialized knowledge and enhance the well-being of the community through cooperation (Durkheim, 1893; Ericsson & Charness, 1994; Lutz & Keil, 2002; Boyd, Richerson, & Henrich, 2011). Knowledge of cause-effect relationships is central to many domains of human cognition and to many types of cultural expertise, from medicine to engineering (Tooby & DeVore, 1987; Wellman & Gelman, 1992; Gopnik & Meltzoff, 1997; Tenenbaum et al., 2007; Goddu & Gopnik, 2024; Keil, 2024). Here we test the hypothesis that people with expert-level causal knowledge automatically activate richer representations in semantic networks of the brain during language comprehension. Specifically, we compare neural responses across physicians and lay people as they read everyday language that elicits implicit causal inferences about illness, such as, “Sam sunbathed on a sunny beach every day. Now he has a cancerous freckle.”

Medical expertise is an ancient and culturally ubiquitous example of human cognitive expertise (Foster, 1976; Ackerknecht, 1982; Norman et al., 2009; Lightner, Heckelsmiller, & Hagen, 2021). In many cultures, medical experts spend a decade or more studying the causes of illness (i.e., etiology, pathophysiology, epidemiology; Schmidt & Boshuizen, 1993; Norman et al., 2009; Nishigori, 2024). The medical expert’s ability to diagnose and treat illness relies on her specialized knowledge of its causes, from evil spirits to energy imbalances and pathogens (Kirmayer, 2004; Lynch & Medin, 2006; Woods, Brooks, & Norman, 2007; Lightner, Heckelsmiller, & Hagen, 2023; de Barra et al., 2025). Such medical expertise is built on a shared foundation of biological knowledge present even in young children, who readily reason about the causes of illness in themselves and others (e.g., *why did my friend get COVID?*; Wellman & Gelman, 1992; Keil, 1992; Inagaki & Hatano, 2006; Legare & Gelman, 2008; Springer & Ruckel, 1992).

Here we tested the hypothesis that even when understanding simple stories that do not require expert-level knowledge, medical experts would show enhanced neural responses to causal inferences about illness in domain-relevant semantic networks (preregistration: https://osf.io/6pnqg). World knowledge is rapidly retrieved during language processing, and prior evidence suggests that individual differences in sensory-motor experience influence neural responses to language (Kutas & Federmeier, 2000; Hagoort et al., 2004; Lau, Phillips, & Poeppel, 2008; Pylkkänen, Oliveri, & Smart, 2009; Yee & Thompson-Schill, 2016; Beilock et al., 2008; Lyons et al., 2010; Wang et al., 2020; Bi, 2021). For example, hockey players exhibit enhanced neural activity in motor networks when listening to action-related language (Beilock et al., 2008; Lyons et al., 2010), and blind and sighted people exhibit partially distinct responses when thinking about the colors of everyday objects presented in words (e.g., “eggplant”) (Wang et al., 2020; Bi, 2021). To our knowledge, how causal expertise impacts the neural basis of language comprehension has not been previously tested.

An alternative, null hypothesis is that when language comprehension does not require expert-level knowledge, experts and non-experts exhibit equivalent neural responses. Prior studies have observed differences between medical experts and novices across large sections of frontal and temporal cortex only in the context of explicit expert-oriented clinical reasoning tasks (Melo et al., 2011, 2017; Durning et al., 2012, 2015; Hruska et al., 2016; Ouellette et al., 2020).

Because illness is exclusive to animate entities, we predicted that medical expertise would enhance responses in temporoparietal semantic regions implicated in the representation of long-term knowledge about animates (i.e., people and animals): the precuneus (PC) and temporoparietal junction (TPJ) (Devlin et al., 2002; Fairhall & Caramazza, 2013a, 2013b; Fairhall et al., 2014; Peer et al., 2015; Wang et al., 2016; Silson et al., 2019; Rabini, Ubaldi, & Fairhall, 2021; Deen & Freiwald, 2025; Aglinskas & Fairhall, 2023; Hauptman, Elli, et al., 2025; Wing et al., 2026). Prior work has found that this temporoparietal network responds to both images and words referring to animate entities (Fairhall & Caramazza, 2013a, 2013b; Fairhall et al., 2014; Wang et al., 2016; Hauptman, Elli, et al., 2025).

Effects of causal expertise could additionally or instead be observed in prefrontal reasoning systems, which are thought to support enhanced reasoning and executive abilities in humans, including causal reasoning (Monti et al., 2007; Duncan, 2010; Donoso, Collins, & Koechlin, 2014; Fugelsang & Dunbar, 2009; Khemlani et al., 2014; Operskalski & Barbey, 2017). Expertise-related changes might also manifest in the frontotemporal language network, which plays a key role in extracting meaning from words and sentences (e.g., Fedorenko et al., 2010; Friederici & Gierhan, 2013). Although some prior studies have reported frontotemporal responses to causal inferences in discourse comprehension, these studies did not localize language, reasoning, or semantic systems in individual participants, leaving open the question of how these different networks contribute to implicit causal inference (Kuperberg et al., 2006; Mason & Just, 2011; Prat et al., 2011).

To test these hypotheses, we asked physicians and age- and education-matched controls to perform a causal inference experiment and a series of localizer tasks, enabling individual-subject localization in each participant. In the main experiment, participants read simple vignettes that invited causal inferences about illness, such as, “Hugh sat by sneezing passengers on the subway. Now he has a case of COVID.” Participants were not told to make explicit causality judgments, instead performing a semantic cover task that required them to attend to the meaning of the text. The content of the vignettes, including their causal connectedness, was accessible to experts and non-experts alike. We hypothesized that the same vignettes would activate richer causal knowledge in medical experts, as evidenced by elevated activity in animacy-related semantic networks of the brain.

We compared neural responses to biological causal inferences about illness to causal inferences about the mechanical breakdown of inanimate objects (e.g., windows, houses) as well as reading causally unconnected sentences that contained illness-related language (e.g., names of illnesses). We hypothesized that expertise effects would be content-specific. In other words, medical expertise should enhance responses to causal inferences about illness in animacy-relevant semantic networks but not responses to causal inferences about inanimate object mechanics in object-relevant semantic networks (Baldassano et al., 2013; Fischer et al., 2016; Silson et al., 2019; Steel et al., 2021; Häusler et al., 2022; Hauptman, Elli, et al., 2025). In a more stringent test of specificity, we hypothesized that medical expertise would enhance responses to causal inferences about biological processes but not mental states, even though both are relevant to animates (Saxe & Kanwisher, 2003; Frith & Frith, 2003; Van Overwalle, 2009; Wolf et al., 2010; Döhnel et al., 2012). A series of localizer tasks was used to identify relevant functional networks – domain-general reasoning, language, and theory of mind – in individual brains (Fedorenko et al., 2010; Monti et al., 2009; Kanjlia et al., 2016; Liu et al., 2020; Saxe & Kanwisher, 2003; Dodell-Feder et al., 2011).

## Method

### Open science practices

The methods and analysis of this experiment were pre-registered prior to data collection (https://osf.io/j76u8). A subset of the current dataset was published in a separate study that used identical experimental procedures and data collection methods (preregistration: https://osf.io/6pnqg; Hauptman & Bedny, 2025).

### Participants

Thirty-two adults (14 women, 18 men, 25-37 years old, *M* = 29.3 years ± 3.5 SD) participated in the study (Supplementary Table 1). Half of the participants (n=16) were physicians employed at Johns Hopkins Medicine (n=15) or surrounding hospitals (n=1). All physicians had completed at least six months of postdoctoral training (i.e., residency, *M* = 2.31 years ± 1.18 SD). The other half of the participants (n=16) were age- and education-matched controls in non-biomedical fields recruited from Johns Hopkins University (n=15) or surrounding universities (n=1). Data from twenty of these participants (n=10 per group) were included in a separate study (Hauptman & Bedny, 2025).

Two additional participants were excluded from the final dataset due to excessive head motion (> 2 mm; n=1) and an image artifact (n=1). One participant included in the final dataset exhibited excessive head motion (> 2 mm) during 1 run of the language/logical reasoning localizer task, which was excluded from analysis.

All participants were screened for cognitive and neurological disabilities (self-report). Participants additionally completed two cognitive tests: the Woodcock-Johnson III (Synonyms, Antonyms, Analogies) and a working memory task (Reading Span; Stone & Towse, 2015). Physicians and non-physicians scored equivalently on both tests (all *p*s > 0.5; Supplementary Figure 1). All participants gave written informed consent and were compensated $30 per hour. The study was reviewed and approved by the Johns Hopkins Medicine Institutional Review Boards (NA_00087983).

### Causal inference experiment

#### Stimuli

Participants read two-sentence vignettes that elicited biological causal inferences about illness, including viral infection, exposure to environmental toxins, genetic factors, and diet (*Illness-Causal* condition; Figure 1B, Supplementary Table 2). The first sentence described something a human agent did or experienced and the second sentence specified the resulting illness (e.g., “Holly used a grimy faucet during her overseas plane ride. Now she has a stomach flu.”). *Illness-Causal* vignettes encoded causal relationships about illness that were accessible to non-physicians, as confirmed by explicit causality judgments collected from a separate group of non-physicians (see Appendix 1 for details). The accessibility of the stimuli allowed us to investigate differences in causal knowledge between the groups as opposed to differences in comprehension. Participants also read two-sentence vignettes that elicited inferences about physical causes of mechanical damage to personally valuable inanimate objects and places (*Mechanical-Causal* condition, e.g., “Haley played an old instrument during her heavy metal show. Now she has a damaged guitar.”). *Mechanical-Causal* vignettes were equally causally related compared to *Illness-Causal* vignettes (Appendix 1) but elicited causal inferences about a different, non-biological domain. The inclusion of this condition enabled us to compare responses to domain-relevant (biological) and non-relevant (non-biological) causal inferences across physicians and non-physicians.

**Figure 1.**
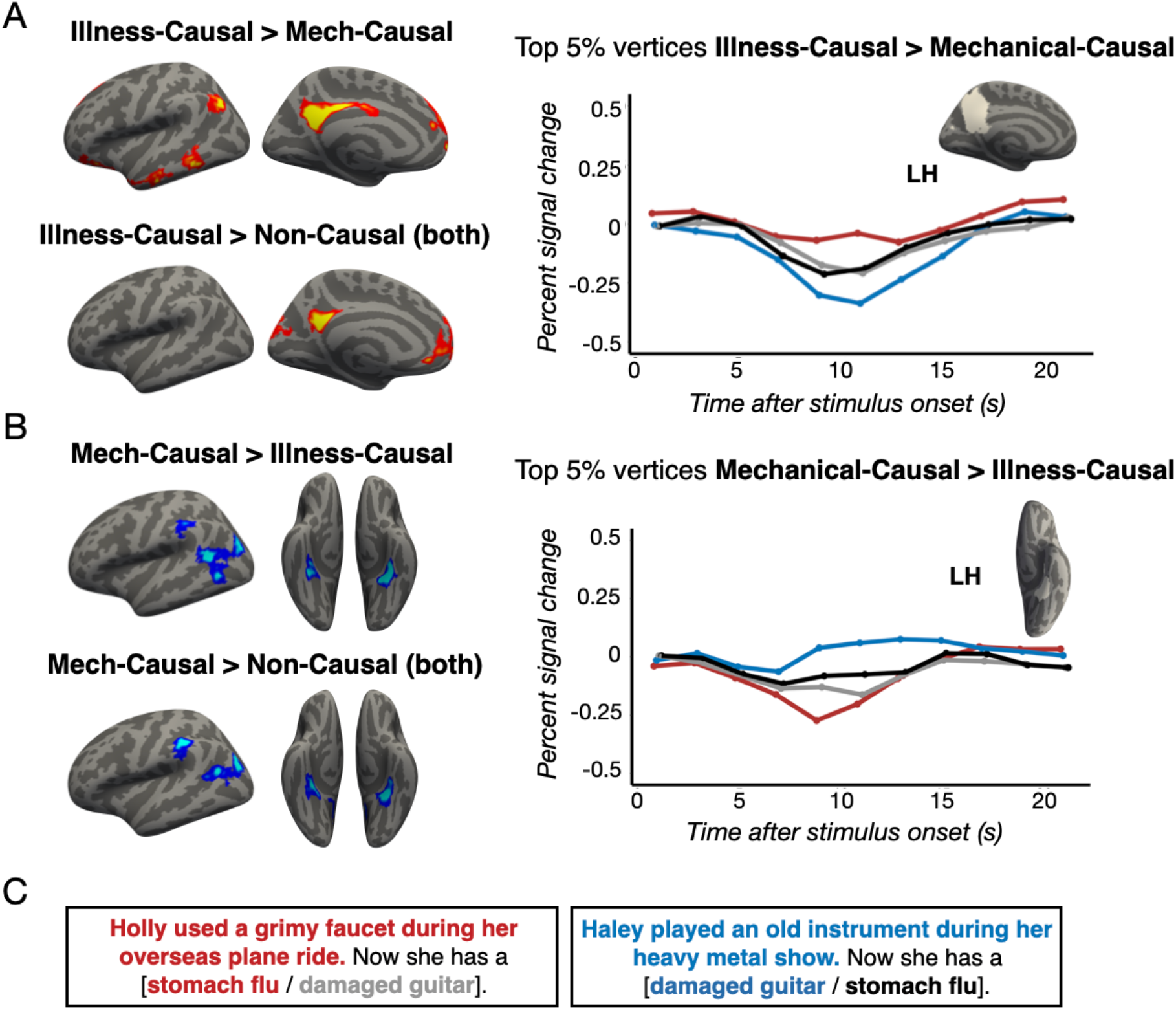
Responses to causal inferences during language comprehension across groups (n=32). Panel A: Causal inferences about illness: whole-cortex results and functional region of interest (fROI) analysis among the top 5% Illness-Causal > Mechanical-Causal vertices in a left PC search space (Dufour et al., 2013). Panel B: Causal inferences about mechanical breakdown: whole-cortex results and fROI analysis among the top 5% Mechanical-Causal > Illness-Causal vertices a in left anterior PPA search space (Hauptman, Elli, et al., 2025). Panel C: Example stimuli. Whole-cortex results are corrected for multiple comparisons (p < .05) and color coded on a scale from p=0.05 to p=0.00001. We hypothesized that the PC and anterior PPA would exhibit opposite domain preferences across groups (see preregistration: https://osf.io/6pnqg).

In addition to reading *Illness-Causal* and *Mechanical-Causal* vignettes, participants read two types of non-causal control vignettes. These vignettes used the same sentences as in the *Illness-Causal* and *Mechanical-Causal* conditions but in a shuffled order: illness cause with mechanical effect (*Noncausal-Illness First*) or mechanical cause with illness effect (*Noncausal-Mechanical First*) (e.g., “Holly used a grimy faucet during her overseas plane ride. Now she has a damaged guitar.” / “Haley played an old instrument during her heavy metal show. Now she has a stomach flu.”). Both causal conditions were rated as more causally related than both non-causal conditions in a separate online experiment (Appendix 1).

All stimuli were matched across multiple linguistic variables, including number of characters, number of words, average number of characters per word, average word frequency, average bigram surprisal (Google Books Ngram Viewer, https://books.google.com/ngrams/), and average syntactic dependency length (Stanford Parser; Marneffe, MacCartney, & Manning, 2006; see Appendix 2 for details).

#### Procedure

We used a “magic detection” task to encourage participants to process the meaning of the vignettes without making explicit causality judgments. Participants saw “magical” catch trials that closely resembled the experimental trials but were fantastical (e.g., “Sadie forgot to wash her face after she ran in the heat. Now she has a cucumber nose.”). On each trial, participants indicated via button press whether “something magical” occurred in the vignette (Yes/No). This semantic cover task encouraged participants to attend to the meaning of the vignettes without revealing the experimental manipulation, as confirmed by a post-participation survey (see Appendix 3 for details).

Both sentences in each vignette were presented simultaneously for 7 s, the first above the second, followed by a 12 s inter-trial interval. The experiment was divided into 6 10-minute runs containing 6-7 trials per condition per run presented in a pseudorandom order (see Appendix 4 for further details about the randomization scheme).

### Language/logical reasoning localizer experiment

To test whether expertise effects manifest in the language and reasoning networks during causal inference elicited by language, we used a localizer task to identify these networks in individual participants. The task had three conditions: language, logical reasoning, and math. In the language condition, participants judged whether two visually presented sentences, one in active and one in passive voice, shared the same meaning. In the logical reasoning condition, participants judged whether two logical statements were consistent (e.g., *If either not Z or not Y then X* vs. *If not X then both Z and Y*). In the math condition, participants judged whether the variable *X* had the same value across two equations (see Liu et al., 2020 for details).

Trials lasted 20 s (1 s fixation + 19 s display of stimuli) and were presented in an event-related design. Participants completed 2 9-minute runs of the task, with trial order counterbalanced across runs and participants. Following prior studies, the language network was identified in individual participants by contrasting *language > math* and the reasoning network by contrasting *logical reasoning > language* (Monti et al., 2009; Kanjlia et al., 2016; Liu et al., 2020).

### Mentalizing localizer experiment

To test whether expertise effects were specific to causal inferences about illness or extended to other animacy-related causal inferences, we used a localizer task to identify neural responses to causal inferences about mental states (i.e., mentalizing; Saxe & Kanwisher, 2003; Dodell-Feder et al., 2011). In this task, participants read 10 mentalizing stories (e.g., a protagonist has a false belief about an object’s location) and 10 physical stories (physical representations depicting outdated scenes, e.g., a photograph showing an object that has since been removed) before answering a true/false comprehension question. We used the mentalizing stories from the original localizer but created stimuli for the physical stories condition that were linguistically matched and incorporated more vivid descriptions of physical interactions (see Hauptman & Bedny, 2025 for details).

Trials were presented in an event-related design, with each one lasting 16 s (12 s stories + 4 s comprehension question) followed by a 12 s inter-trial interval. Participants completed 2 5-minute runs of the task, with trial order counterbalanced across runs and participants. Responses to mentalizing were identified in individual participants by contrasting *mentalizing stories* > *physical stories* (Saxe & Kanwisher, 2003; Dodell-Feder et al., 2011).

### Data acquisition

Whole-brain fMRI data was acquired at the F.M. Kirby Research Center of Functional Brain Imaging on a 3T Phillips Achieva Multix X-Series scanner. T1-weighted structural images were collected in 150 axial slices with 1 mm isotropic voxels using the magnetization-prepared rapid gradient-echo (MP-RAGE) sequence. Functional T2*-weighted BOLD scans were collected using a gradient echo planar imaging (EPI) sequence with the following parameters: 36 sequential ascending axial slices, repetition time (TR) = 2 s, echo time (TE) = 0.03 s, flip angle = 70°, field of view (FOV) matrix = 76 × 70, slice thickness = 2.5 mm, inter-slice gap = 0.5, slice-coverage FH = 107.5, voxel size = 2.4 × 2.4 × 3 mm, PE direction = L/R, first order shimming. Data were acquired in one experimental session lasting approximately 120 minutes. All stimuli were visually presented on a rear projection screen with a Cambridge Research Systems BOLDscreen 32 UHD LCD display (image resolution = 1920 x 1080) using custom scripts written in PsychoPy3 (https://www.psychopy.org/, Peirce et al., 2019). Participants viewed the screen via a front-silvered, 45° inclined mirror attached to the top of the head coil.

### fMRI data preprocessing and general linear model (GLM) analysis

Preprocessing included motion correction, high-pass filtering (128 s), mapping to the cortical surface (Freesurfer), spatially smoothing on the surface (6 mm FWHM Gaussian kernel), and prewhitening to remove temporal autocorrelation. Covariates of no interest included signal from white matter, cerebral spinal fluid, and motion spikes.

For the causal inference experiment, the main GLM modeled the four main conditions (*Illness-Causal*, *Mechanical-Causal*, *Noncausal-Illness First, Noncausal-Mechanical First*) and the “magical” catch trials during the 7 s display of the vignettes after convolving with a canonical hemodynamic response function and its first temporal derivative. This GLM additionally included participant response time and the number of human agents described in each vignette (one agent or more than one) as covariates of no interest. In a separate GLM used for an exploratory analysis, we modeled each individual trial in each run. For the mentalizing localizer experiment, a separate predictor was included for each condition (*mentalizing stories, physical stories*), modeling the 16 s display of each story and corresponding comprehension question. For the language/logical reasoning localizer experiment, a separate predictor was included for each of the three conditions (*language, logical reasoning, math*), modeling the 20 s duration of each trial.

Runs were modeled separately and combined within-subject using a fixed-effects model (Dale, Fischl, & Sereno, 1999; Smith et al., 2004). Group-level random-effects analyses were corrected for multiple comparisons across the whole cortex at *p* < .05 family-wise error rate (FWER) using a nonparametric permutation test (cluster-forming threshold *p* < .01 uncorrected) (Winkler et al., 2014; Eklund, Nichols, & Knutsson, 2016; Eklund, Knutsson, & Nichols, 2019).

### Univariate individual-subject fROI analysis

We defined individual-subject functional ROIs (fROIs) in animacy regions (precuneus (PC), temporoparietal junction (TPJ), and fusiform face area (FFA)), an “inanimates” region (anterior parahippocampal place area (PPA)), and the language and logical reasoning networks. For all analyses, percent signal change (PSC) was extracted and averaged over the entire duration of the trial (17 s total), starting at 4 s to account for hemodynamic lag.

Our main question was whether physicians would show enhanced responses to causal inferences about illness in a temporoparietal semantic network previously implicated in knowledge about animates (e.g., PC, TPJ). To this end, we created illness inference fROIs in bilateral PC and TPJ group search spaces (Dufour et al., 2013) using an iterated leave-one-run-out procedure, which allowed us to perform sensitive individual-subjects analysis while avoiding statistical non-independence (Vul & Kanwisher, 2011). In each participant, we identified the most responsive vertices to illness causal inferences in bilateral PC and TPJ search spaces separately in 5 of the 6 runs (top 5% of vertices, *Illness-Causal* > *Mechanical-Causal*). We extracted PSC for each condition compared to rest in the held-out run, averaging the results across all iterations. We then compared the difference between *Illness-Causal* and *Mechanical-Causal* across participant groups (physicians, non-physicians) using repeated-measures ANOVAs.

We used a similar approach to conduct several control analyses. First, we tested whether any group differences in responses to causal inferences about illness emerged outside the temporoparietal semantic network implicated in thinking about animates, i.e., in the language and logical reasoning networks. Language fROIs were identified by selecting the most language-responsive vertices (top 5%) in left frontal and temporal language areas (search spaces: Fedorenko et al., 2010) using the *language* > *math* contrast from the language/logical reasoning localizer. Reasoning fROIs were identified by selecting the most logic-responsive vertices (top 5%) in a left frontoparietal network (search space: Liu et al., 2020) using the *logical reasoning > language* contrast. In each fROI, we extracted PSC for all conditions in the illness causal inference task and compared the difference between *Illness-Causal* and *Mechanical-Causal* across groups. We used the same approach to test for sensitivity to causal inferences about illness across groups in the fusiform face area (FFA) (search space: Julian et al., 2012). Prior work suggests that the FFA responds to images of animates (i.e., faces) but is not involved in high-level semantics (e.g., Kanwisher et al., 1997; Grill-Spector et al., 2004; Konkle & Caramazza, 2013; Fairhall & Caramazza, 2013a; Fairhall et al., 2014; Wang et al., 2016; Hauptman, Elli, et al., 2025). We therefore predicted that this region would not exhibit sensitivity to animacy-related causal inferences elicited by language.

To test whether any group differences in the temporoparietal network were limited to inferences about illness or extended to other animacy-related causal inferences (i.e., mentalizing), we created mentalizing fROIs in left PC and TPJ (search spaces: Dufour et al., 2013; top 5% of vertices, *mentalizing stories > physical stories*, leave-one-run-out approach). Lastly, we created mechanical inference fROIs in a network previously implicated in thinking about inanimate objects (i.e., left anterior PPA search spaces; Hauptman, Elli, et al., 2025; top 5% of vertices, *Mechanical-Causal* > *Illness-Causal*, leave-one-run-out approach). This analysis enabled us to test whether group differences also emerged for mechanical causal inferences and were thus not tied to a particular semantic domain. These analyses focused on the left hemisphere, where group differences in responses to causal inferences about illness were observed.

### Searchlight MVPA

We used a linear support vector machine classifier (PyMVPA toolbox; Hanke et al., 2009) to compare decoding between illness causal inferences and mechanical causal inferences (*Illness-Causal* vs. *Mechanical-Causal*) in physicians and non-physicians across the whole cortex using a 10 mm radius spherical searchlight (according to geodesic distance, to better respect cortical anatomy over Euclidean distance; Glasser et al., 2013). This yielded for each participant a classification map indicating the classifier’s accuracy in the neighborhood surrounding every vertex. Individual subject searchlight accuracy maps were averaged within group (physicians, non-physicians). The non-physicians map was then subtracted from the physicians map to identify the regions where decoding was better in the physician group. We used a permutation-based correction approach to threshold this difference map by randomly shuffling the individuals assigned to each group and performing the same subtraction of averaged group maps 1000 times. This yielded an empirical null distribution of subtracted values for each vertex that was used to threshold the true group difference map at *p* < .05.

### Within-vs. across-group correlations of responses to causal inferences about illness

In an exploratory analysis, we tested whether responses to illness causal inferences (*Illness-Causal*) in the “animacy semantic network” (i.e., PC, TPJ) were more similar within participant groups compared to across groups. Higher within-group neural similarity would suggest that medical training (or lack thereof) shapes neural responses to causal inferences about illness in similar ways. Following the results of univariate fROI analysis, we focused on the neural similarity of illness inferences in the left PC and left TPJ.

First, we identified the top 300 vertices most responsive to illness causal inferences (*Illness-Causal > Mechanical-Causal*) in left PC and TPJ group search spaces (Dufour et al., 2013) in individual participants. We then obtained one observation (z-scored beta parameter estimate of the individual-trial GLM) per trial of the *Illness-Causal* condition (n=38 trials). To reduce noise, we averaged data from pairs of trials with similar content (see Appendix 5 for details). We then calculated the ranked similarity (Euclidean distance) between each pair of observations. In fROIs that exhibited average cross-participant correlations greater than zero (assessed via one-tailed Student’s *t*-tests), we subsequently performed a more sensitive within- vs. across-group analysis. For each participant in a group, we correlated (Spearman’s rho) their ranked similarity values for illness causal inference trials with i) the average ranked similarities of the remaining participants in that group and ii) the average ranked similarity values of all participants in the other group.

This yielded two correlation values (within-group and across-group) for each participant. We then performed a repeated-measures ANOVA to test for differences in within- vs. across-group similarity in both physicians and non-physicians.

## Results

### Behavioral results

For the main causal inference experiment, participants’ accuracy was near ceiling on the task across all conditions (physicians: *M* = 98.1% ± 2.38 SD, non-physicians: *M* = 97.9% ± 2.43 SD). “Magical” trials were dropped from remaining analyses. There were no differences in accuracy or response time across groups in the “magic detection” task (Supplementary Figure 2; repeated measures ANOVA, condition (*Illness-Causal*, *Mechanical-Causal*, *Noncausal-Illness First*, *Noncausal-Mechanical First*) x group (physicians, non-physicians), accuracy: main effect of group, *F*_(1,30)_ = 0, *p* = 1, group x condition interaction, *F*_(3,90)_ = 2.03, *p* = .12; response time: main effect of group, *F*_(1,30)_ = 0, *p* = .99; group x condition interaction, *F*_(3,90)_ = 0.38, *p* = .77). Across both groups, participants were less accurate in their judgments about *Noncausal-Illness First* vignettes compared to all other conditions in the causal inference task (i.e., *Illness-Causal*, *Mechanical-Causal*, and *Noncausal-Mechanical First*; main effect of condition, *F*_(3,90)_ = 4.24, *p* = .007). Additionally, participants’ responses were faster in their judgments about *Illness-Causal* vignettes compared to both non-causal conditions (*Noncausal-Illness First*, *Noncausal-Mechanical First*; main effect of condition, *F*_(3,90)_ = 45.33, *p* < .001). Response time was included as a covariate of no interest in the GLM.

For the localizer tasks, there were likewise no group differences (all *p*s > 0.25), except for a marginal group difference in response time in the language/logical reasoning localizer task, where non-physicians responded faster than physicians in *logical reasoning* trials (repeated measures ANOVA, condition (*language*, *logical reasoning*, *math*) x group (physicians, non-physicians), group x condition interaction, *F*_(2,60)_ = 3.06, *p* = .05) (see Appendix 6 for details).

### Neural signatures of causal expertise in a temporoparietal “animacy semantic network”

#### ROI analysis

Inferring the causes of illness during language comprehension elicited left-lateralized neural responses in a temporoparietal semantic network previously implicated in knowledge about animate entities (e.g., people, animals). Precuneus (PC) and temporoparietal junction (TPJ) fROIs exhibited increased activity to causal inferences about illness compared to mechanical causal inferences (individual-subject fROI leave-one-run-out analysis; repeated measures ANOVA, condition (*Illness-Causal*, *Mechanical-Causal*) x hemisphere (left, right) x group (physicians, non-physicians), PC: main effect of condition, *F*_(1,30)_ = 27.45, *p* < .001, TPJ: main effect of condition, *F*_(1,30)_ = 11.41, *p* = .002). The difference between illness and mechanical causal vignettes was more pronounced in the left hemisphere (PC: main effect of hemisphere, *F*_(1,30)_ = 0.25, *p* = .62, condition x hemisphere interaction, *F*_(1,30)_ = 39.97, *p* < .001, TPJ: main effect of hemisphere, *F*_(1,30)_ = 4.78, *p* = .04, condition x hemisphere interaction, *F*_(1,30)_ = 9.10, *p* = .005). By contrast, sensitivity to causal inferences about illness was not observed in the fusiform face area (FFA), which has been shown to respond to images of animates (e.g., Kanwisher et al., 1997; Konkle & Caramazza, 2013; see Appendix 7 for details).

Critically, temporoparietal animacy regions (i.e., the PC and TPJ) also showed effects of expertise. In both regions, there was a condition x hemisphere x group interaction (PC: *F*_(1,30)_ = 6.42, *p* = .02; TPJ: *F*_(1,30)_ = 5.92, *p* = .02; see Appendix 8 for full results). Post-hoc analyses revealed that physicians showed higher responses to illness causal inferences than mechanical causal inferences in both the left PC and left TPJ (repeated measures ANOVA, condition (*Illness-Causal*, *Mechanical-Causal*) x group (physicians, non-physicians), left PC: condition x group interaction, *F*_(1,30)_ = 4.56, *p* = .04; right PC: condition x group interaction, *F*_(1,30)_ = 0.36, *p* = .55; left TPJ: condition x group interaction, *F*_(1,30)_ = 10.66, *p* = .003; right TPJ: condition x group interaction, *F*_(1,30)_ = 0.72, *p* = .40; Figure 2; Appendix 8).

**Figure 2.**
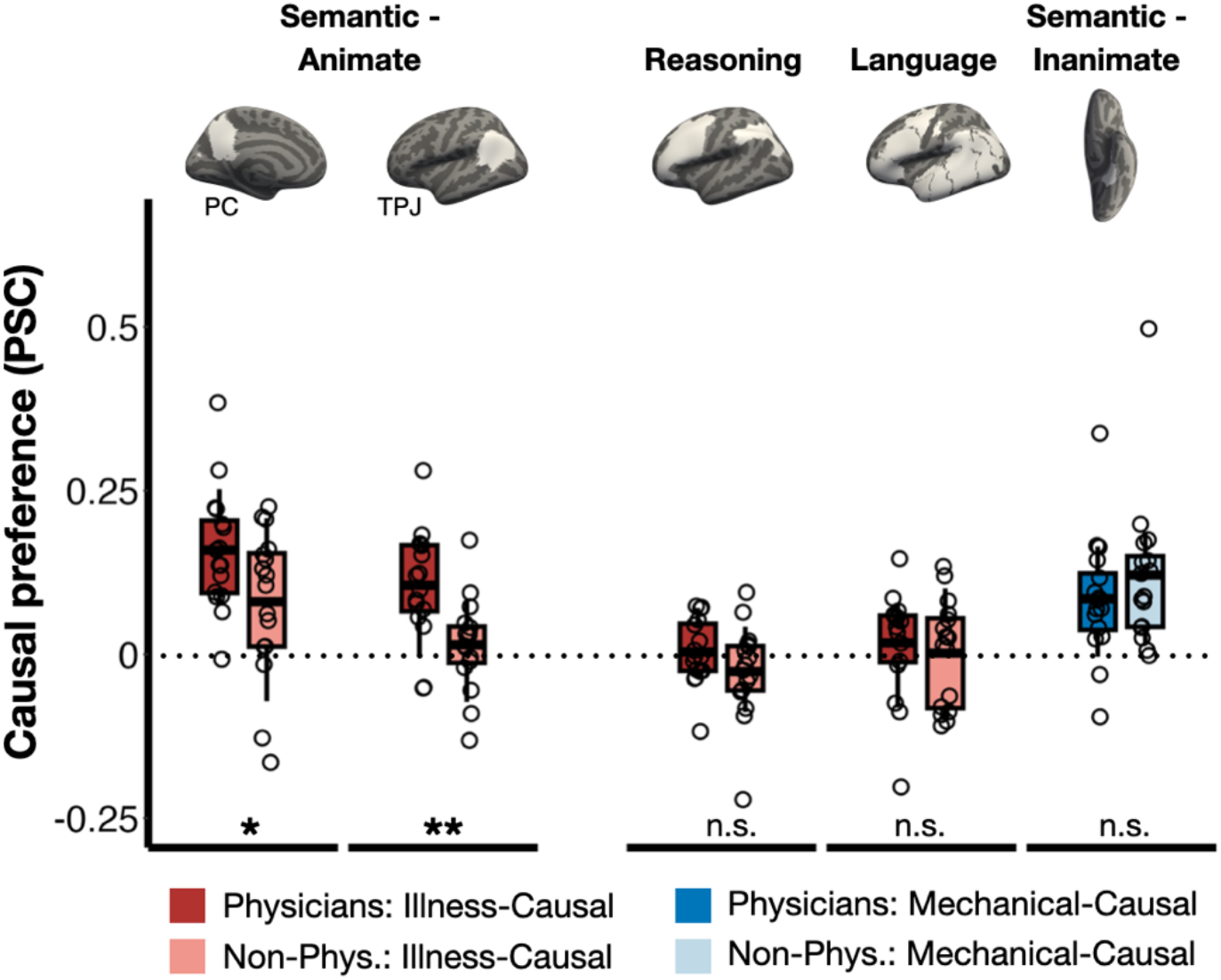
Enhanced responses to causal inferences about illness in animacy semantic network of medical experts, individual-subject fROI analysis. Differences in average percent signal change (PSC) between causal conditions (Illness-Causal > Mechanical-Causal in red, Mechanical-Causal > Illness-Causal in blue; top 5% of vertices, leave-one-run-out analyses). The dotted horizontal line indicates 0 (i.e., no difference between conditions). PC: left precuneus, TPJ: left temporoparietal junction, Reasoning: left logical reasoning network, Language: left language network, Semantic - Inanimate: left anterior parahippocampal place area. Significance codes: ‘***’ 0.001 ‘**’ 0.01 ‘*’ 0.05 ‘.’ 0.1 ‘n.s.’ 1.

Across groups, causal inferences about illness also elicited higher responses than non-causal conditions in the PC. Only physicians showed this effect in the TPJ (see Appendix 9 for full results).

#### Whole-cortex analysis

In whole-cortex analysis of the full sample (n=32), causal inferences about illness elicited elevated neural responses in a temporoparietal semantic network implicated in thinking about animates (i.e., people and animals) compared to mechanical causal and non-causal vignettes (i.e., bilateral PC, with larger effects on the left, and left TPJ; *p* < .05, corrected for multiple comparisons; Figure 1A, Supplementary Figures 3-4). Responses to causal inferences about illness in the PC were spatially adjacent to but distinct from responses to mental state inferences (i.e., mentalizing) (Figure 3B, Appendix 10; Saxe & Kanwisher, 2003). Conversely, a semantic network previously implicated in thinking about inanimate objects and intuitive physical reasoning (i.e., bilateral anterior parahippocampal regions, left supramarginal gyrus, and left posterior superior temporal sulcus) was more active for mechanical causal inferences compared to illness causal and non-causal vignettes (*p* < .05, corrected for multiple comparisons, Figure 1B, Supplementary Figures 3-4).

**Figure 3.**
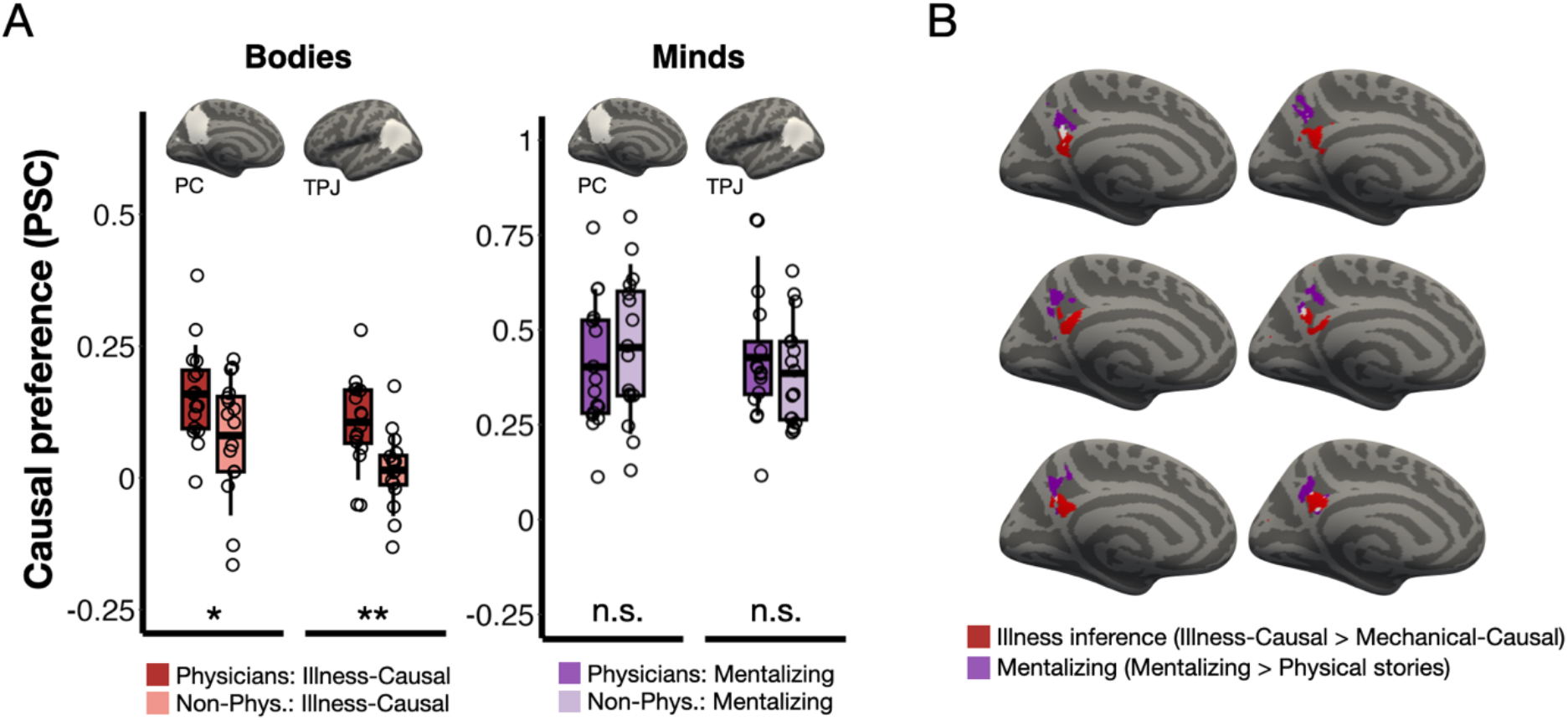
Medical expertise responses are specific to inferences about bodies as opposed to minds. Panel A: Group differences in univariate responses to causal inferences about illness (Bodies, red) mentalizing (Minds, purple), individual-subject fROI analysis. Panel B: Spatial dissociation between univariate responses to illness inferences and mental state inferences in the PC.

When comparing groups in whole-cortex univariate analysis, we found that no vertices reached significance. However, in multivariate whole-cortex searchlight analysis, the PC and TPJ exhibited higher decoding accuracy between illness and mechanical causal inferences in physicians than non-physicians. Group differences in decoding accuracy were also observed in left medial prefrontal cortex, bilateral insula, and right middle and inferior temporal gyrus (Supplementary Figure 6). In addition, subject-wise overlap maps for the *Illness-Causal > Mechanical-Causal* contrast showed more robust and/or spatially consistent responses to illness causal inferences in physicians’ bilateral PC and left TPJ (Figure 4A).

**Figure 4.**
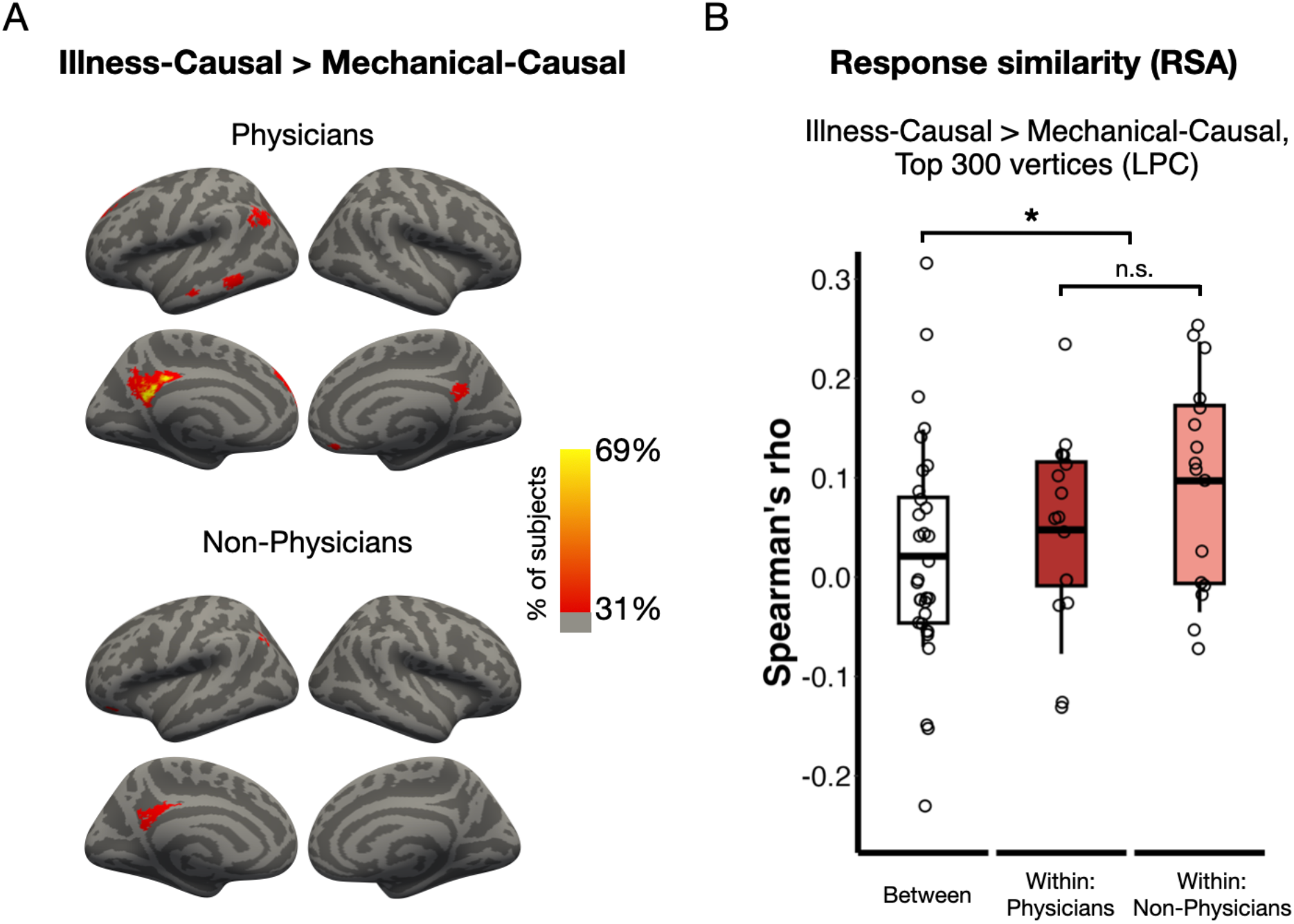
Medical experts exhibit more robust and self-similar responses to causal inferences about illness. Panel A: Across-subject overlap of responses to causal inferences about illness compared to causal inferences about mechanical breakdown within each group. Vertices are color-coded according to the proportion of participants who showed activation (one-tailed, p < .05 uncorrected, cluster correction 50 mm^2^). Panel B: Within- and between-group similarity in precuneus (PC) responses to causal inferences about illness across trials. Significance codes: ‘***’ 0.001 ‘**’ 0.01 ‘*’ 0.05 ‘.’ 0.1 ‘n.s.’ 1.

### No effects of medical expertise in frontoparietal reasoning/executive regions or frontotemporal language regions

Enhanced responses to causal inferences about illness were specific to the temporoparietal semantic network implicated in thinking about animates and were not found in reasoning, language, or other semantic networks. No group differences were observed in the frontoparietal reasoning or frontotemporal language networks (individual-subject fROI analysis; Figure 2; repeated measures ANOVA, condition (*Illness-Causal*, *Mechanical-Causal*) x group (physicians, non-physicians), reasoning network, combined frontal and parietal search spaces: main effect of group, *F*_(1,30)_ = 1.68, *p* = .21, main effect of condition, *F*_(1,30)_ = 0.77, *p* = .39, condition x group interaction, *F*_(1,30)_ = 1.35, *p* = .25; language network, combined frontal and temporal search spaces: main effect of group, *F*_(1,30)_ = 0.13, *p* = .73, main effect of condition, *F*_(1,30)_ = 0.45, *p* = .51, condition x group interaction, *F*_(1,30)_ = 0.29, *p* = .59).

There were also no effects of expertise in semantic networks responsive to inanimate entities (i.e., the anterior PPA; Figure 2; individual-subject fROI leave-one-run-out analysis; repeated measures ANOVA, condition (*Mechanical-Causal*, *Illness-Causal*) x hemisphere (left, right) x group (physicians, non-physicians), main effect of condition, *F*_(1,30)_ = 31.12, *p* < .001, hemisphere x group interaction, *F*_(1,30)_ = 0, *p* = .94, condition x group interaction, *F*_(1,30)_ = 1.76, *p* = .19, condition x hemisphere x group interaction, *F*_(1,30)_ = 0.02, *p* = .9; see Appendix 11 for full results).

### Medical expertise effects are specific to causal inferences about illness (bodies) rather than mental states (minds)

In an exploratory analysis, we tested whether medical expertise enhances responses to all inferences about people, including inferences about their minds, or only responses to inferences about the body (biology). To distinguish between these hypotheses, we compared responses to mental state inferences during a mentalizing localizer task across physicians and non-physicians (Saxe & Kanwisher, 2003; Dodell-Feder et al., 2011; Hauptman & Bedny, 2025). There were no differences between physicians and non-physicians in the magnitude of responses to mental state inferences in the left PC or TPJ, with equally robust responses observed across groups (Figure 3A; repeated measures ANOVA, condition (*mentalizing stories*, *physical stories*) x group (physicians, non-physicians), left PC: main effect of condition, *F*_(1,29)_ = 192.57, *p* < .001, main effect of group, *F*_(1,29)_ = 1.57, *p* = .22, condition x group interaction, *F*_(1,29)_ = 1.16, *p* = .29; left TPJ: main effect of condition, *F*_(1,29)_ = 207.4, *p* < .001, main effect of group, *F*_(1,29)_ = 0.13, *p* = .72, condition x group interaction, *F*_(1,29)_ = 0.08, *p* = .79). These results suggest that medical expertise specifically enhances neural responses to causal inferences about illness as opposed to all animacy-related inferences.

### Physicians show more similar neural responses to each other than to non-physicians

In an exploratory analysis, we compared the similarity of neural responses to causal inferences about illness across trials in physicians and non-physicians. We reasoned that physicians and non-physicians might have slightly but measurably different interpretations of the illness causal vignettes. If so, we would expect to observe more coherent neural responses to causal inferences about illness among physicians than across physicians and non-physicians. First, we investigated whether either left PC or left TPJ showed consistent spatial patterns of activity across all participants during causal inferences about illness. The left PC exhibited consistent patterns across the full sample (*t*_(31)_ = 3.01, *p* = .005), but the left TPJ did not (*t*_(31)_ = 0.8, *p* = .43). We therefore pursued a comparison of within- vs. across-group correlations only in the left PC.

Both physicians and non-physicians showed higher correlations within-group compared to across-groups (Figure 4B; repeated measures ANOVA, correlation type (within-group, across-group) x group (physicians, non-physicians): main effect of within vs. across group correlation, *F*_(1,30)_ = 5.23, *p* = .03, main effect of group, *F*_(1,30)_ = 0.76, *p* = .39, within/between x group interaction, *F*_(1,30)_ = 0.99, *p* = .33). Thus, when making causal inferences about illness, physicians’ neural responses were more similar to other physicians’ responses than to non-physicians’ responses, and the opposite was true for non-physicians.

## Discussion

Compared to people without medical training, physicians show enhanced neural responses when making implicit causal inferences about illness from language. Effects of medical expertise are specific to the temporoparietal semantic network previously implicated in representing knowledge about animate entities (i.e., people, animals): the left precuneus (PC) and left temporoparietal junction (TPJ) (Devlin et al., 2002; Fairhall & Caramazza, 2013a, 2013b; Fairhall et al., 2014; Peer et al., 2015; Wang et al., 2016; Silson et al., 2019; Rabini, Ubaldi, & Fairhall, 2021; Deen & Freiwald, 2025; Aglinskas & Fairhall, 2023; Hauptman, Elli, et al., 2025). The neural specificity of this expertise effect and equivalent responses and response times across groups on the “magic detection task” together suggest that the observed group differences are unlikely to be related to faster prediction or greater attention to illness-related language in physicians. Instead, we hypothesize that the observed neural differences between groups reflect differences in the richness of the causal knowledge activated by medical causal experts compared to non-experts. These findings suggest that an animacy-responsive temporoparietal semantic network undergoes experience-based functional plasticity in people who acquire expert-level causal knowledge about biology. Alternatively, people with enhanced neural responses to biological causal knowledge may “self-select” to become physicians. Future studies comparing neural responses to causal knowledge about illness in the same individuals before and after completing medical training can distinguish between these possibilities.

Differences between physicians and non-physicians were layered on top of common cross-group sensitivity to causal inferences. Across groups, causal inferences about illness activated a temporoparietal network previously shown to respond to both images and words referring to animate entities (i.e., PC, TPJ; Fairhall & Caramazza, 2013a, Silson et al., 2019; Deen & Freiwald, 2025; Wing et al., 2026). This result is in line with prior work using naturalistic stimuli that implicates these semantic networks in the construction of causal situation models based on incoming discourse (Johnson-Laird, 1993; Zwaan & Radvansky, 1998; Baldassano et al., 2017; Lee & Chen, 2022). By contrast, causal inferences about mechanical breakdown activated a network that responds to words and images referring to inanimate large objects and places (i.e., anterior PPA; Baldassano et al., 2013; Fairhall et al., 2014; Silson et al., 2019; Steel et al., 2021; Häusler et al., 2022; Hauptman, Elli, et al., 2025). Together, these findings suggest that causal knowledge retrieved during language comprehension is embedded in distributed content-specific semantic networks.

Across both physicians and non-physicians, neural responses to causal inferences about illness were anatomically specific within the broader animacy network. Lateral ventral occipitotemporal cortex (i.e., the fusiform face area), which responds to visual images of animate entities, did not show sensitivity to causal inferences about illness (e.g., Kanwisher et al., 1997; Konkle & Caramazza, 2013). Within temporoparietal cortex, responses to causal inferences about illness were anatomically adjacent to but distinct from responses to mental state inferences (i.e., mentalizing) (Saxe & Kanwisher, 2003; Frith & Frith, 2003; Van Overwalle, 2009; Wolf et al., 2010; Döhnel et al., 2012; see also Hauptman & Bedny, 2025). The proximity of these neural responses is intriguing given some prior developmental work suggesting that children’s early understanding of biology is initially rooted in their understanding of human minds (e.g., plants grow because people want them to) (Carey, 1985). Other work suggests that young children recognize the distinct “causal laws” that govern psychological vs. biological processes (Springer & Keil, 1991; Wellman & Gelman, 1992; Inagaki & Hatano, 1993, 2004; Keil, 1994). Examining the development of the neural distinction between causal inferences about the body vs. the mind is a promising avenue for future research.

Unlike semantic networks, the frontotemporal language network did not show enhanced responses to causal inferences or effects of expertise despite being highly activated during story reading. This result offers additional evidence against the possibility that the group differences observed in the current study are due to physicians’ increased attention to illness-related stimuli. Prior work finds that the language network is responsive to semantic information at the level of individual words, phrases, and sentences, suggesting that this network builds “shallow” representations of linguistic input that are subsequently sent to other functional networks for deeper processing (Pallier, Devauchelle, & Dehaene, 2011; Blank & Fedorenko, 2020; Shain et al., 2024; Casto et al., 2025). Our results are consistent with this hypothesis. Although language is a major channel by which we convey causal information about the world (e.g., Wolff, 2003; Harris & Koenig, 2006; Lombrozo & Carey, 2006; Bar-Asher Siegal, 2026), such information does not appear to be represented in the language network itself. If the primary role of the language network is to support communication rather than thought, it may be robust to individual variation in semantic knowledge (Carruthers, 2002; Pinker & Jackendoff, 2005; Gleitman & Papafragou, 2005; Fedorenko, Piantadosi, & Gibson, 2024).

Likewise, we did not observe expertise effects or enhanced responses to implicit causal inferences in frontoparietal reasoning systems. Unlike semantic networks, the frontoparietal reasoning network showed higher activity during *non-causal* compared to causal vignettes. This result suggests that unlike explicit causal reasoning, implicit causal inferences that occur spontaneously during language comprehension do not engage domain-general reasoning systems and may be more akin to automatic semantic retrieval than effortful goal-directed reasoning (e.g., Swinney et al., 1979; Swaab et al., 1998).

In sum, although multiple neural systems are engaged when understanding discourse, semantic networks appear to play a special role in enabling implicit causal inference and are a primary target of functional change in experts who acquire specialized causal knowledge.

An open question for future research is whether cross-cultural and developmental differences in biological knowledge show similar neural signatures to those of biomedical expertise. The current study compares neural responses to causal inferences about illness in individuals with and without formal biomedical training. Individual differences in biological knowledge are also present across cultures and development. For instance, intuitive theories of biology across cultures posit diverse underlying causal mechanisms for illness, including energy imbalances, disruptions in “vital force,” and encounters with magical or spiritual beings (Foster, 1976; Ackerknecht, 1982; Kirmayer, 2004; Patwardhan et al., 2005; Legare & Gelman, 2008; Legare & Shtulman, 2017; Lightner, Heckelsmiller, & Hagen, 2023; Lynch & Medin, 2006; de Barra et al., 2025). Additionally, within the same culture, children’s causal biological knowledge changes substantially over the course of development (Carey, 1985; Simons & Keil, 1995; Au & Romo, 1999; Inagaki & Hatano, 2004; Bares & Gelman, 2008). Future work can explore how such cultural and developmental differences manifest in semantic networks of the brain during implicit causal inference.

To conclude, we find that the neural basis of automatic causal inferences elicited during language comprehension is shaped by cultural expertise, even when comprehension does not require expert-level knowledge. Content-specific semantic networks of the brain encode causal knowledge retrieved during language comprehension and are modified by causal expertise.

## Supplementary Figures

**Supplementary Figure 1.**
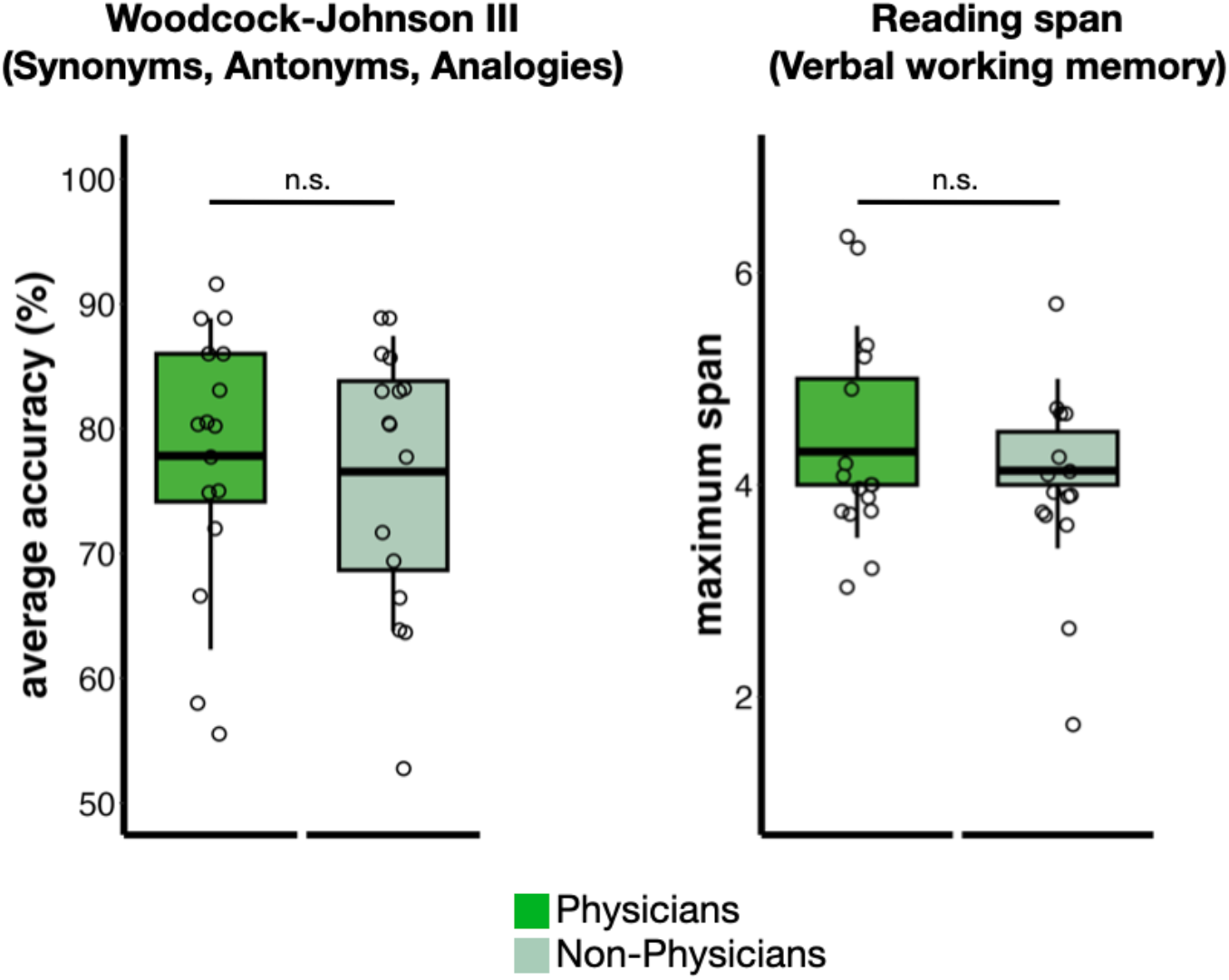
Cognitive test results by group. The horizontal line within each boxplot indicates the mean. Physician and non-physician participants scored equivalently on both the Woodcock-Johnson III (Synonyms, Antonyms, Analogies combined score: *t*_(30)_ = 0.34, *p* = .74) and a working memory task (maximum numbers recalled correctly: *t*_(29)_ = 0.56, *p* = .58). Working memory task results from one non-physician participant were not included due to experimenter error.

**Supplementary Figure 2.**
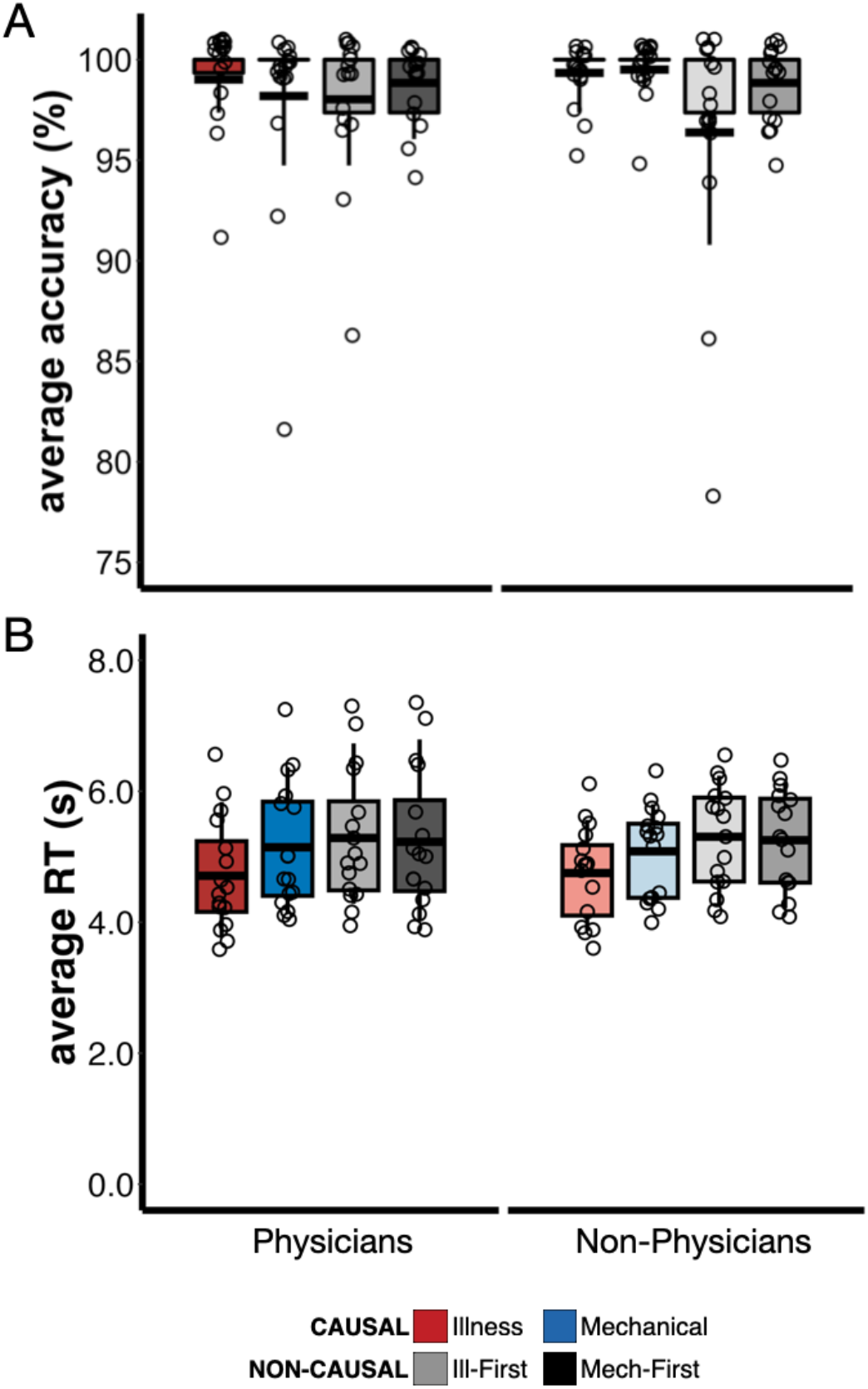
Behavioral results from the illness inference task across groups. The horizontal line within each boxplot indicates the mean. There were no group differences in accuracy or response time across conditions (see Appendix 6 for full behavioral results). Response time was entered as a covariate of no interest in the GLM.

**Supplementary Figure 3.**
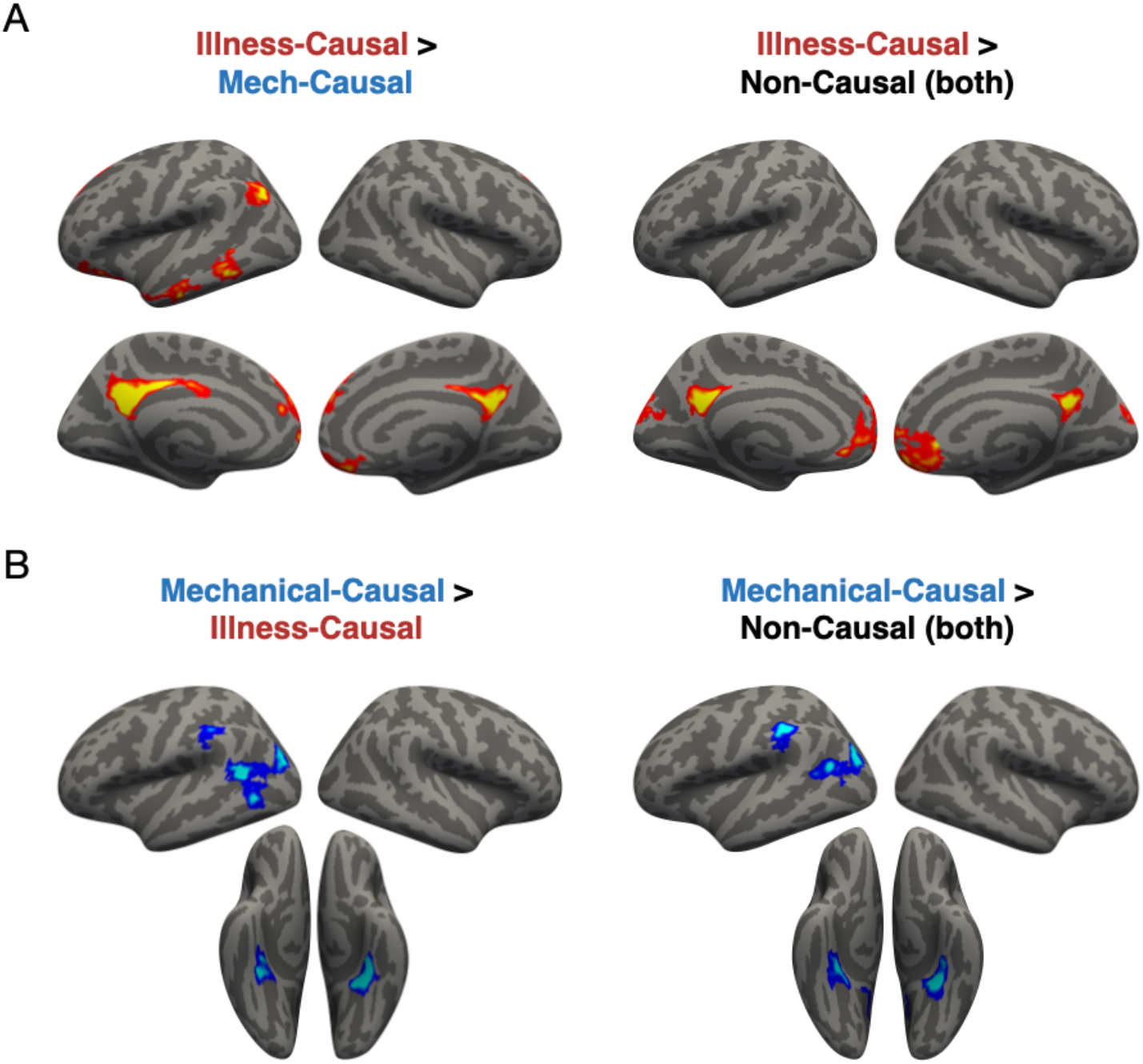
Whole-cortex results (one-tailed) for causal vs. non-causal vignettes, combined across groups (n=32). Panel A: *Illness-Causal* > *Mechanical-Causal* and *Illness-Causal* > *Non-Causal* (both versions of non-causal vignettes combined), corrected for multiple comparisons (*p* < .05 FWER permutation corrected, cluster-forming threshold *p* < .01 uncorrected). Panel A: *Mechanical-Causal* > *Illness-Causal* and *Mechanical-Causal* > *Non-Causal* (both versions of non-causal vignettes), corrected for multiple comparisons (*p* < .05 FWER permutation corrected, cluster-forming threshold *p* < .01 uncorrected). All vertices are color coded on a scale from p=0.05 to p=0.00001.

**Supplementary Figure 4.**
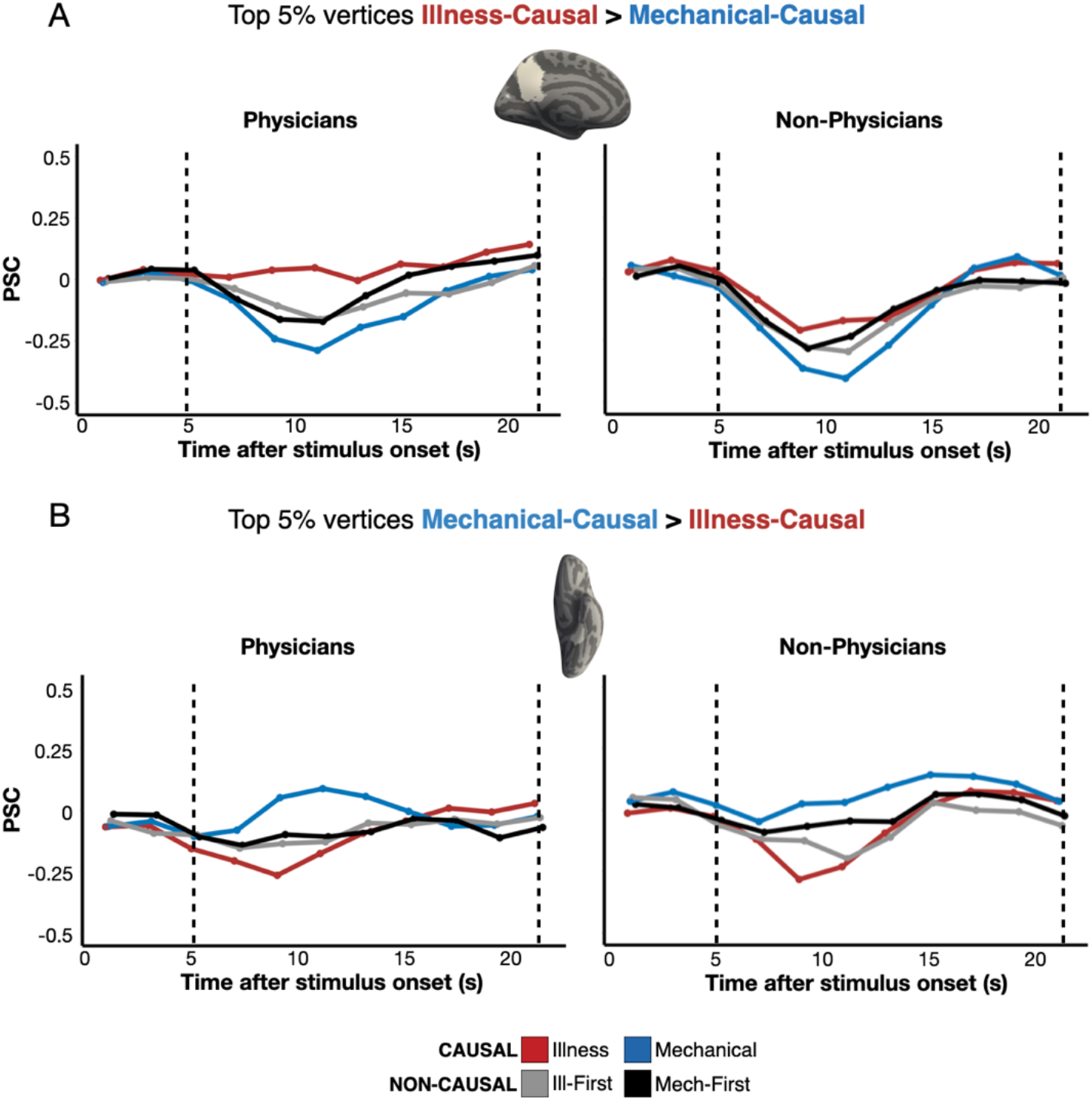
Individual-subject fROI analysis of responses to causal inferences about illness and causal inferences about mechanical breakdown across groups. Panel A: Percent signal change (PSC) for each condition among the top 5% *Illness-Causal* > *Mechanical-Causal* vertices in a left PC search space (Dufour et al., 2013) in individual participants, established via a leave-one-run-out analysis. Panel B: PSC for each condition among the top 5% *Mechanical-Causal* > *Illness-Causal* vertices in a left anterior PPA search space (Hauptman, Elli, et al., 2025) in individual participants, established via a leave-one-run-out analysis.

**Supplementary Figure 5.**
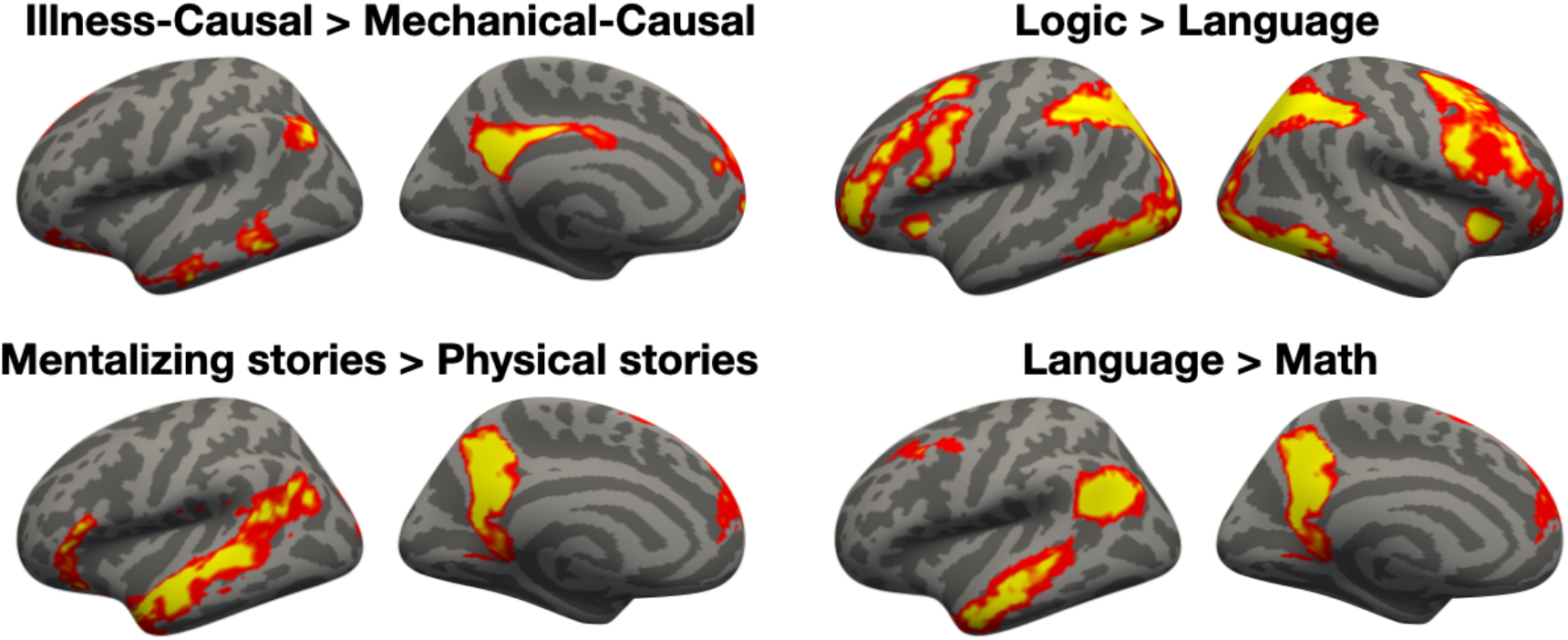
Functional localization of mentalizing, language, and logical reasoning networks and comparison to responses to causal inferences about illness (Monti et al., 2009; Fedorenko et al., 2010; Dodell-Feder et al., 2011; Liu et al., 2020). Group maps for each contrast of interest (n=32, one-tailed) are corrected for multiple comparisons (*p* < .05 FWER, cluster-forming threshold *p* < .01 uncorrected). Vertices are color coded on a scale from p=0.05 to p=0.00001 (Illness-Causal > Mechanical-Causal) and from p=0.001 to p=0.0000001 (localizer contrasts).

**Supplementary Figure 6.**
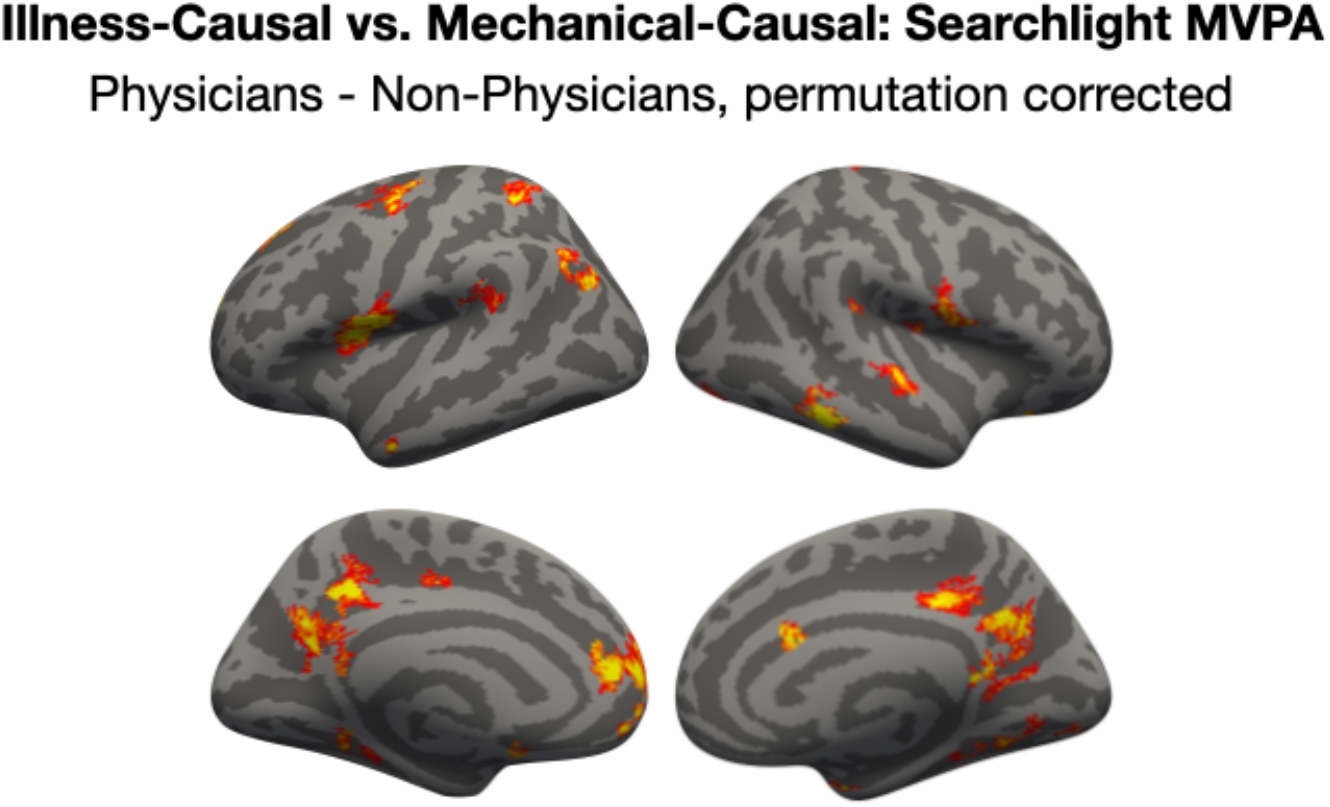
Differences in decoding accuracy (*Illness-Causal* vs. *Mechanical-Causal*) between physicians and non-physicians. Whole-brain searchlight maps were averaged within-group, subtracted from each other (physicians minus non-physicians), and thresholded based on a permutation correction procedure (see Appendix 12). Vertices are color coded on a scale from 5-15% (difference in decoding accuracy between groups).

**Supplementary Figure 7.**
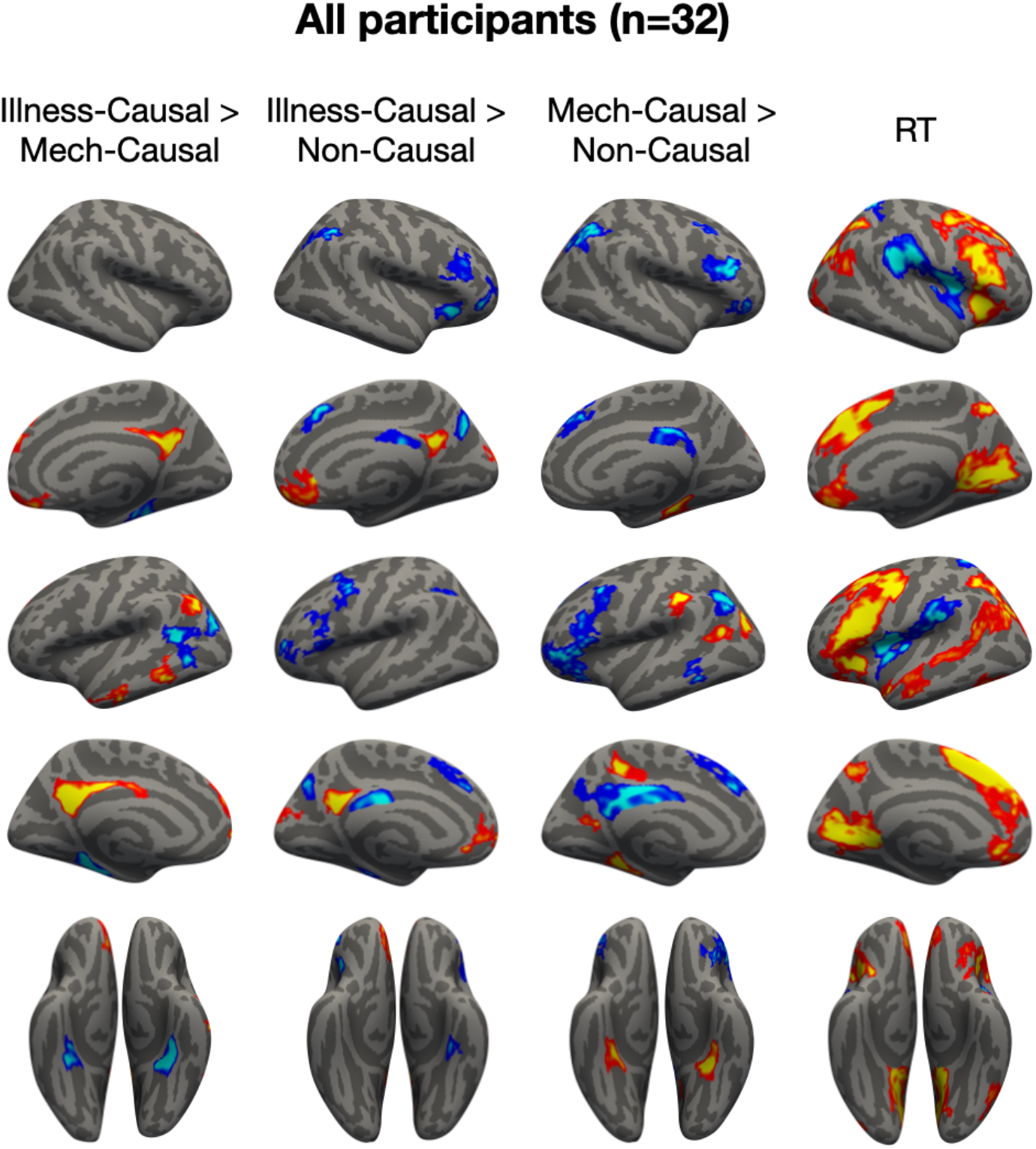
Full whole-cortex univariate results from combined participant groups (n=32). Regions whose activity scales with response time (RT) are displayed under “RT.” Group maps (two-tailed) are corrected for multiple comparisons (*p* < .05 FWER, cluster-forming threshold *p* < .01 uncorrected). Vertices are color coded on a scale from p=0.05 to p=0.00001.

**Supplementary Figure 8.**
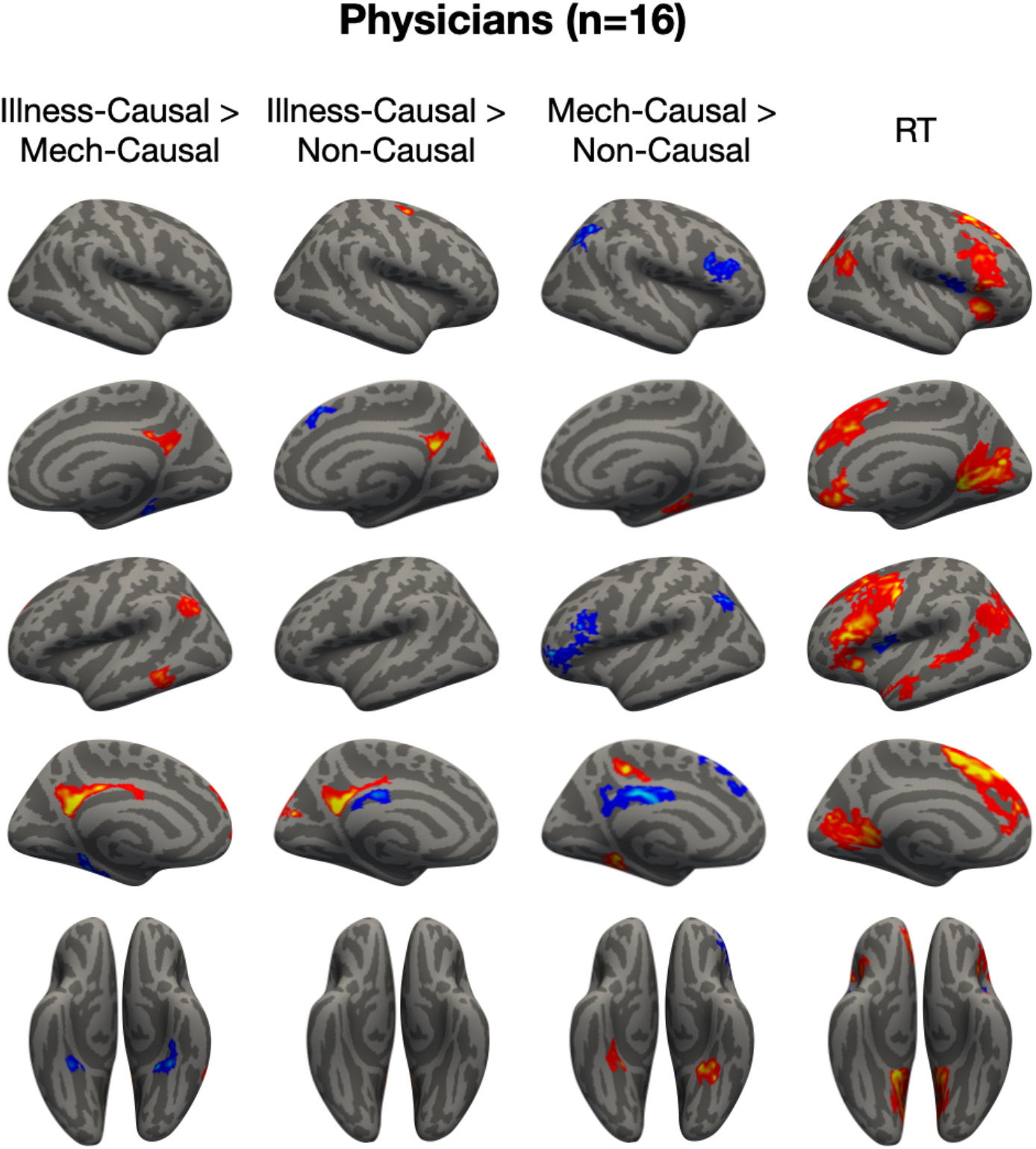
Full whole-cortex univariate results from the physician group (n=16). Regions whose activity scales with response time (RT) are displayed under “RT.” Group maps (two-tailed) are corrected for multiple comparisons (*p* < .05 FWER, cluster-forming threshold *p* < .01 uncorrected). Vertices are color coded on a scale from p=0.05 to p=0.00001.

**Supplementary Figure 9.**
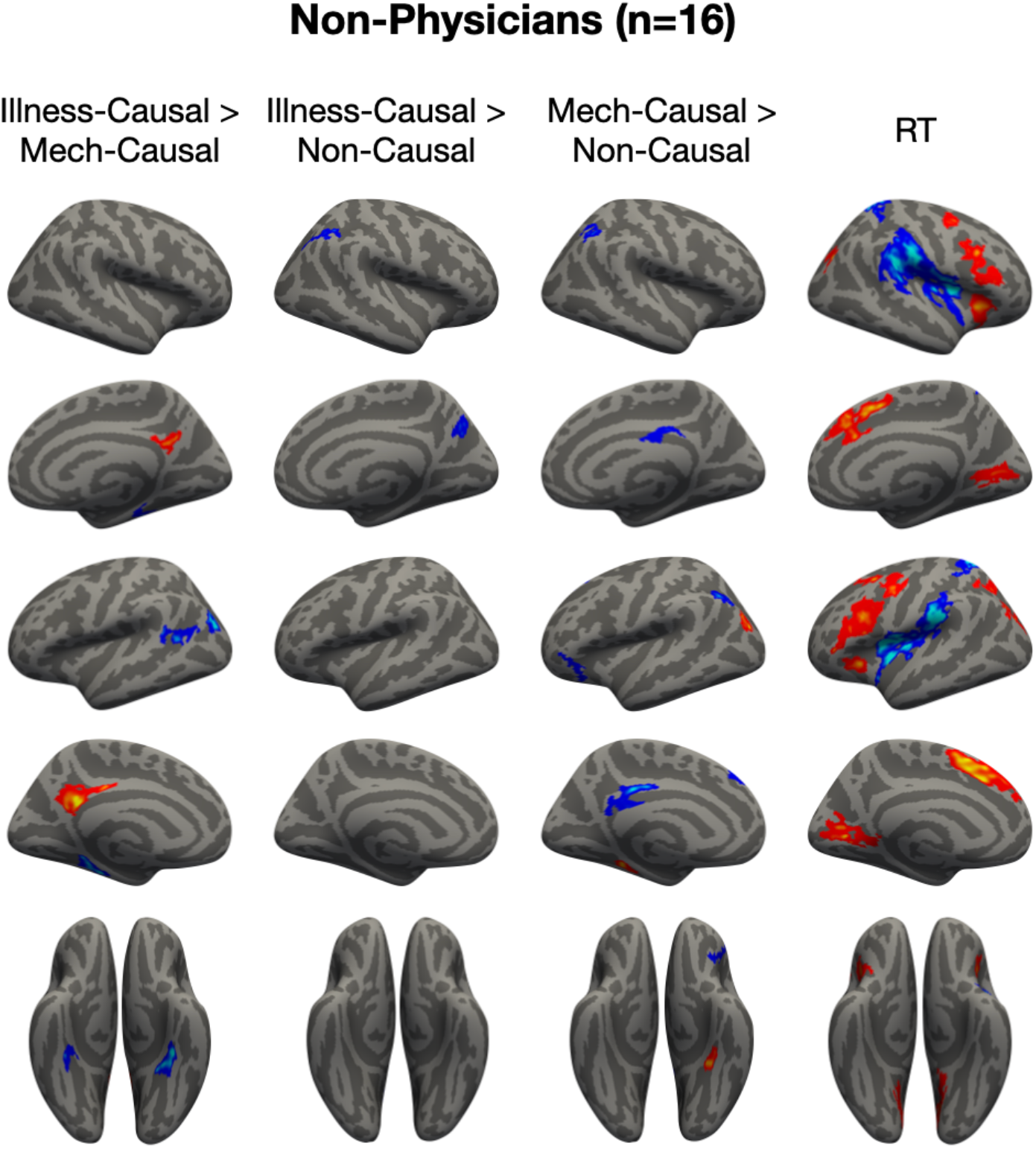
Full whole-cortex univariate results from the non-physician group (n=16). Regions whose activity scales with response time (RT) are displayed under “RT.” Group maps (two-tailed) are corrected for multiple comparisons (*p* < .05 FWER, cluster-forming threshold *p* < .01 uncorrected). Vertices are color coded on a scale from p=0.05 to p=0.00001.

**Supplementary Figure 10.**
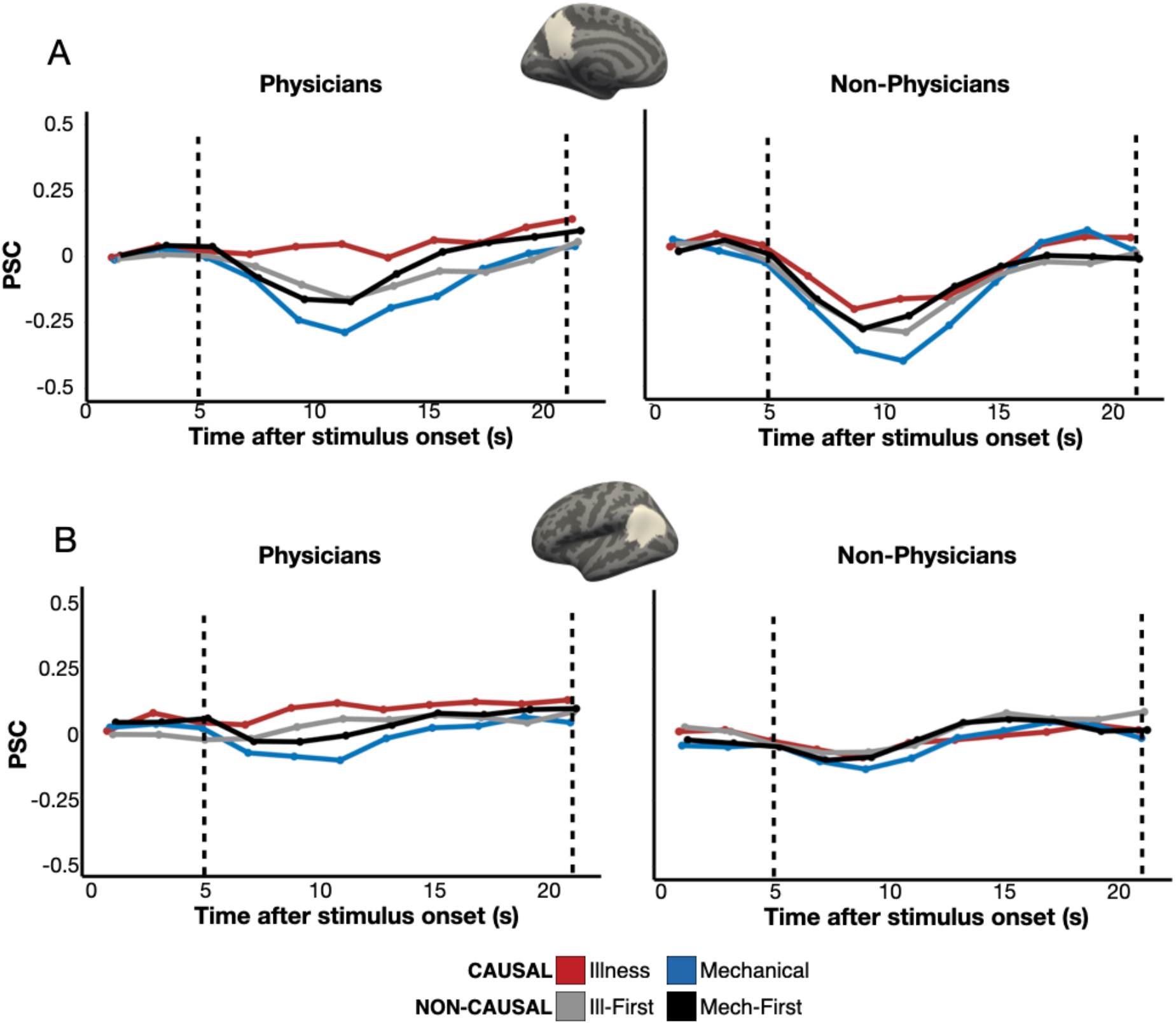
Time-courses of responses to illness inferences in PC and TPJ across groups. Percent signal change (PSC) for each condition among the top 5% *Illness-Causal* > *Mechanical-Causal* vertices in left PC and TPJ search spaces (Dufour et al., 2013) in individual participants, established via a leave-one-run-out analysis, is shown. An illustration of the group differences in responses in the critical window (5-21 s, marked by dotted lines) is shown in Figure 2. Participant dispersion data are shown in Supplementary Figure 11.

**Supplementary Figure 11.**
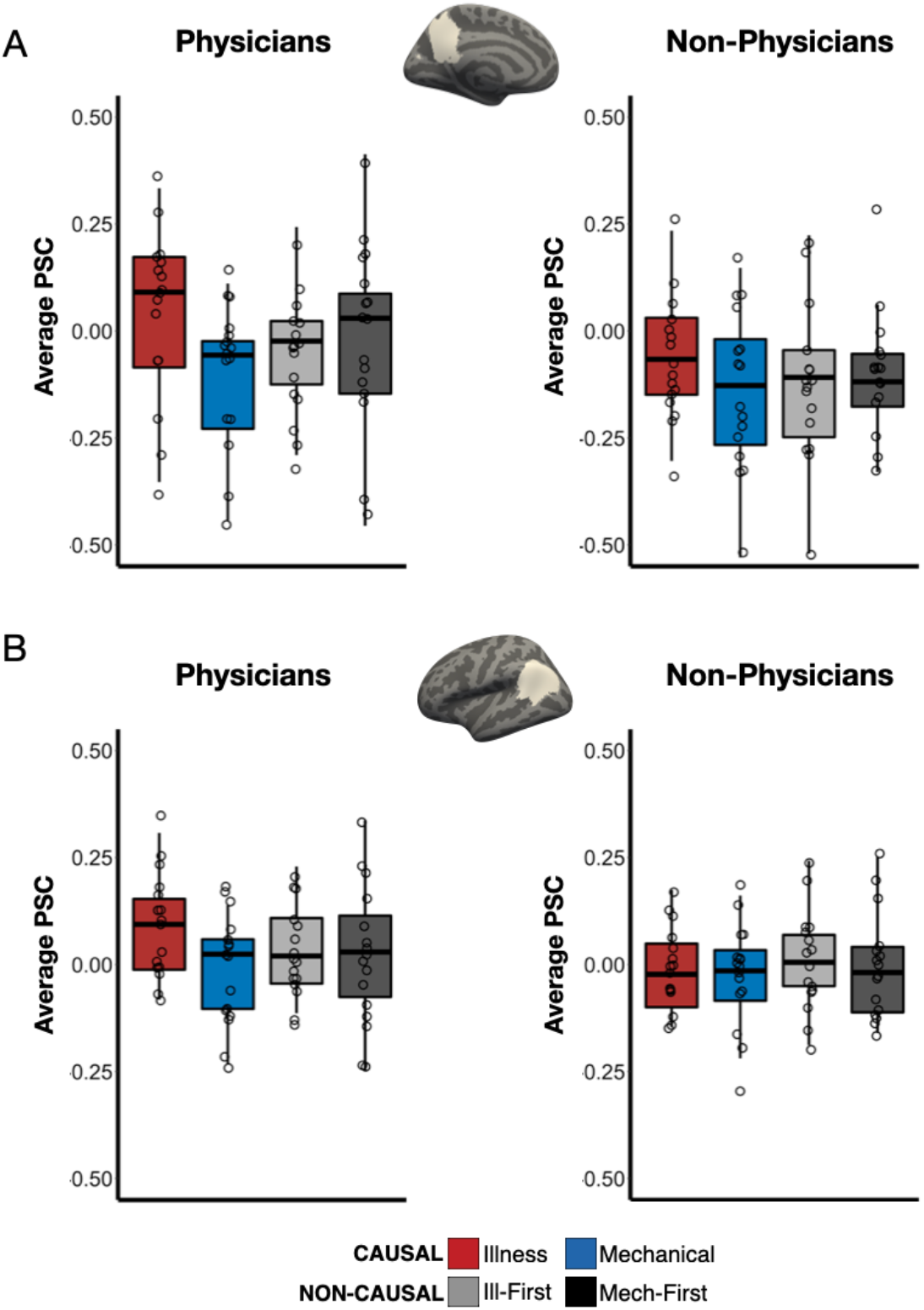
Participant dispersion data of average PSC in top 5% *Illness-Causal > Mechanical-Causal* vertices in left PC and TPJ across groups (see Figure 2 for group differences, Supplementary Figure 10 for time-courses). Boxplots display average PSC in the critical window (5-21 s) across groups and fROIs. The horizontal line within each boxplot indicates the mean.

**Supplementary Figure 12.**
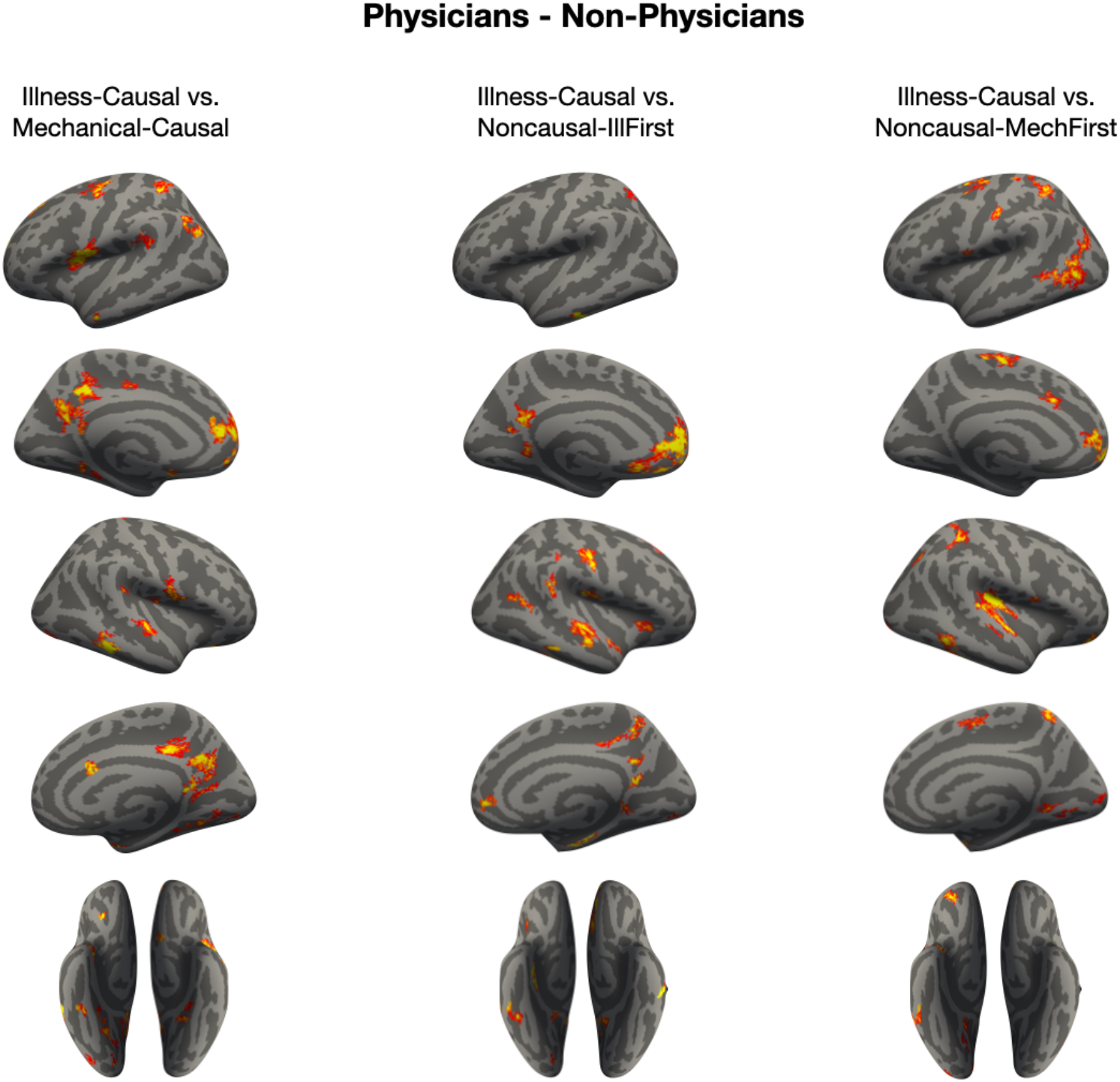
Full results: differences in decoding accuracy between physicians and non-physicians. Whole-brain searchlight maps were averaged within-group, subtracted from each other (physicians minus non-physicians), and thresholded based on a permutation correction procedure (see Appendix 12). Vertices are color coded on a scale from 5-15% (difference in decoding accuracy between groups).

## Supplementary Tables

**Supplementary Table 1.**
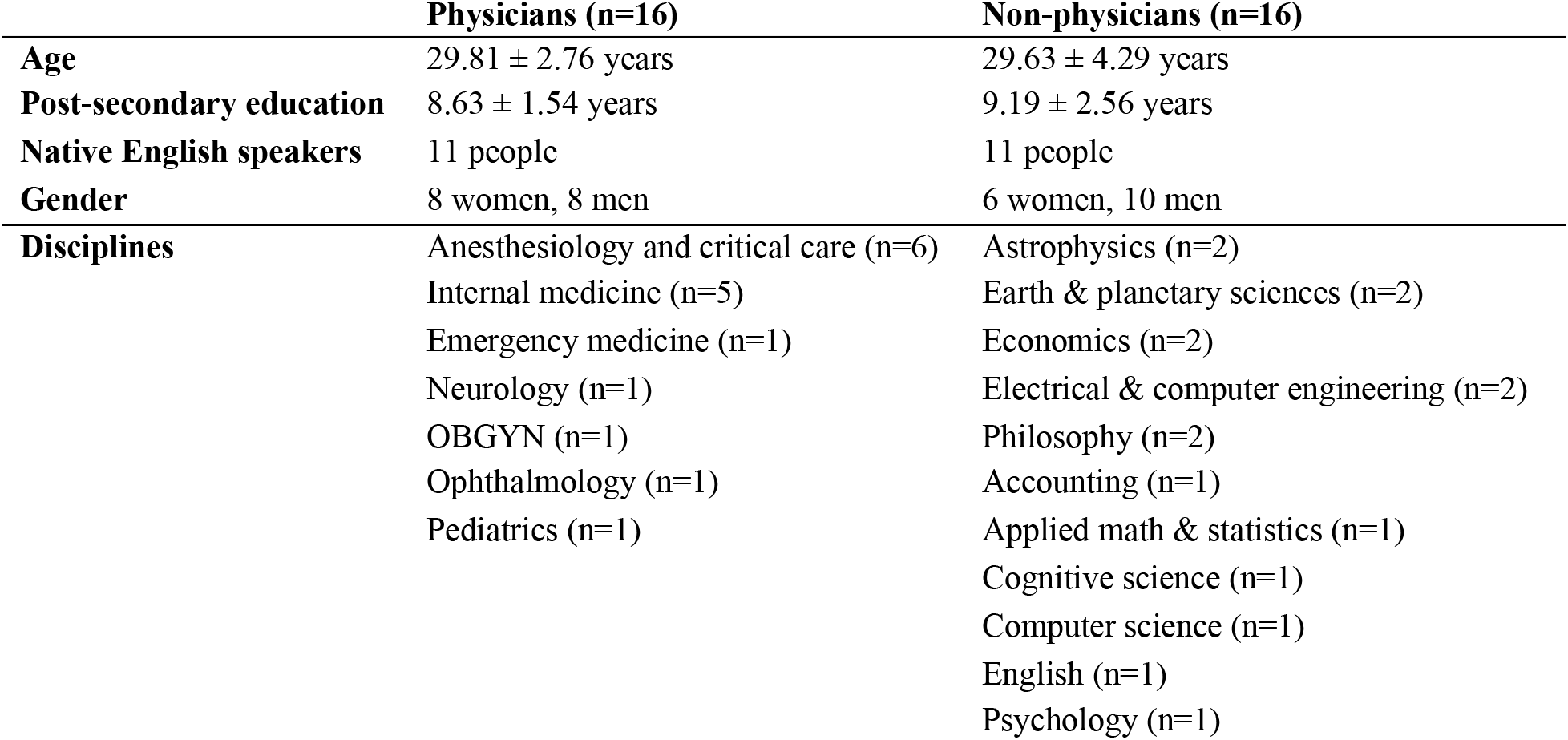
Participant information.

|  | **Physicians (n=16)** | **Non-physicians (n=16)** |
|---|---|---|
| **Age** | 29.81 ± 2.76 years | 29.63 ± 4.29 years |
| **Post-secondary education** | 8.63 ± 1.54 years | 9.19 ± 2.56 years |
| **Native English speakers** | 11 people | 11 people |
| **Gender** | 8 women, 8 men | 6 women, 10 men |
| **Disciplines** | Anesthesiology and critical care (n=6) | Astrophysics (n=2) |
|  | Internal medicine (n=5) | Earth & planetary sciences (n=2) |
|  | Emergency medicine (n=1) | Economics (n=2) |
|  | Neurology (n=1) | Electrical & computer engineering (n=2) |
|  | OBGYN (n=1) | Philosophy (n=2) |
|  | Ophthalmology (n=1) | Accounting (n=1) |
|  | Pediatrics (n=1) | Applied math & statistics (n=1) |
|  |  | Cognitive science (n=1) |
|  |  | Computer science (n=1) |
|  |  | English (n=1) |
|  |  | Psychology (n=1) |

**Supplementary Table 2:** Types of illnesses and their causes in the illness causal inference condition (*Illness-Causal*). Each row corresponds to a single *Illness-Causal* vignette (total n=38) seen by each participant. Vignettes assigned the same “pair number” were averaged for the purposes of an exploratory similarity analysis. A list of the vignettes can be found on OSF (https://osf.io/j76u8).

| **Illness type** | **Cause** | **Pair number** |
|---|---|---|
| COVID | viral infection | 1 |
| COVID | viral infection | 1 |
| HIV-AIDS | viral infection | 2 |
| malaria | viral infection | 2 |
| chickenpox | viral infection | 3 |
| flu | viral infection | 3 |
| flu | viral infection | 4 |
| flu | viral infection | 4 |
| food poisoning | bacterial infection | 5 |
| stomach flu | viral infection | 5 |
| cold | viral infection | 6 |
| cold | viral infection | 6 |
| acne | exposure to irritants | 7 |
| acne | exposure to irritants | 7 |
| lung disease | exposure to toxins | 8 |
| lung disease | exposure to toxins | 8 |
| brain cancer | exposure to toxins | 9 |
| throat cancer | exposure to toxins | 9 |
| pneumonia | exposure to toxins | 10 |
| allergic reaction | exposure to allergens | 10 |
| GI inflammation | exposure to toxins | 11 |
| GI inflammation | exposure to toxins | 11 |
| lung cancer | exposure to toxins | 12 |
| blood cancer | exposure to toxins | 12 |
| liver disease | alcoholism | 13 |
| heart disease | diet | 13 |
| high blood pressure | diet | 14 |
| type 2 diabetes | diet | 14 |
| anemia | diet | 14 |
| breast cancer | genetic factors | 15 |
| cancer (unspecified) | genetic factors | 15 |
| heart disease | genetic factors | 16 |
| heart disease | genetic factors | 16 |
| liver disease | genetic factors | 17 |
| epilepsy | genetic factors | 17 |
| skin cancer | sun exposure | 18 |
| skin cancer | sun exposure | 18 |
| asthma | prematurity | N/A |

**Supplementary Table 3:** Results of preregistered MVPA for *Illness-Causal* vs. *Mechanical-Causal* in individual-subject functional ROIs (see Appendix 12 for details). Each fROI was created by selecting the top 300 vertices for each contrast (see “Contrast”) in a PC search space (Dufour et al., 2013). Accuracy refers to classifier performance against chance (50%) for *Illness-Causal* vs. *Mechanical-Causal*. Permuted and Bonferroni-corrected (across fROIs) p-values are reported. Ment_vs_phys: *mentalizing stories* > *physical stories* (mentalizing localizer). Caus_vs_rest: *Illness-Causal + Illness-Mechanical* > *Rest.* Phys.: physician; Non-phys.: non-physician.

| **Search space** | **Contrast** | **Group** | **Accuracy** | **t** | **Permuted p** | **Bonferroni adj. p** |
|---|---|---|---|---|---|---|
| LPC | ment_vs_phys | phys. | 69.30% | 9.77 | 0 | 0 |
| LPC | ment_vs_phys | non-phys. | 54.70% | 1.21 | 0.1355 | 0.4884 |
| LPC | caus_vs_rest | phys. | 63% | 3.74 | 0.0025 | 0.004 |
| LPC | caus_vs_rest | non-phys. | 58.30% | 1.82 | 0.0509 | 0.1768 |

## Appendices

## Appendix 1 Online experiment protocol

Prior to the fMRI experiment, we collected explicit causality judgments from a separate group of online participants (n=30). Each online participant read all vignettes from the causal inference experiment (152 vignettes) in addition to 12 filler vignettes that were designed to be either maximally causally related or unrelated (164 vignettes total), one vignette at a time. Their task was to judge the extent to which it was possible that the event described in the first sentence of each vignette caused the event described in the second sentence on a 4-point scale (1 = not possible; 4 = very possible). 4 participants were excluded on the basis of inaccurate responses on the filler trials (i.e., difference between average ratings for maximally causally related and maximally causally unrelated vignettes <2). Among the 26 remaining participants, 12 read vignettes from Version A and 14 read vignettes from Version B of the experiment. To eliminate erroneous responses, we first excluded trials with RTs 2.5 SD outside their respective condition means within participants and then excluded trials with outlier RTs (more than 1.5 IQR below Q1 or more than 1.5 IQR above Q3) across participants (approximately 5% of all trials excluded in total). We found that both causal conditions (*Illness-Causal*, *Mechanical-Causal*) were more causally connected than both non-causal conditions, *t*(25) = 36.97, *p* < .001 (causal: *M* = 3.51 ± 0.78 SD, non-causal: *M* = 1.10 ± 0.45 SD). In addition, *Illness-Causal* and *Mechanical-Causal* items received equally high causality ratings, *t*(25) = −0.64, *p* = .53 (*Illness-Causal*: *M* = 3.49 ± 0.77 SD, *Mechanical-Causal*: *M* = 3.53 ± 0.79 SD).

## Appendix 2 Details on stimulus creation

*Illness-Causal* and *Mechanical-Causal* vignettes were constructed in pairs, such that each member of a given pair shared parallel or near-parallel phrase structure. All conditions were matched (pairwise t-tests, all *p*s > 0.3, no statistical correction) on multiple linguistic variables known to modulate neural activity in language regions (e.g., Pallier, Devauchelle, & Dehaene, 2011; Shain, Blank et al., 2020). These included number of characters, number of words, average number of characters per word, average word frequency, average bigram surprisal (Google Books Ngram Viewer, https://books.google.com/ngrams/), and average syntactic dependency length (Stanford Parser; Marneffe, MacCartney, & Manning, 2006). All conditions were matched for all linguistic variables across the first sentence, second sentence, and the entire vignette.

Sentences that were incorrectly parsed by the automatic syntactic parser (i.e., past participle adjectives parsed as verbs) were corrected by hand. Word frequency was calculated as the negative log of a word’s occurrence rate in the Google corpus between the years 2017-2019. Bigram surprisal was calculated as the negative log of the frequency of a given two-word phrase in the Google corpus divided by the frequency of the first word of the phrase. This calculation uses a log base of 2 in order to express surprisal in terms of “bits” that the first word provides in the context of the phrase. We used *bigram* surprisal as our surprisal measure to maximize the number of n-grams that had an entry in the corpus. Even so, 64 out of the 1515 total bigrams (4%) did not have an entry in the corpus and were therefore assigned the highest surprisal value among the rest of the bigrams (see Willems et al., 2016).

## Appendix 3 Details on participant recruitment and post-participation survey

Physician participants were recruited via email advertisements distributed by chief residents and Johns Hopkins Medicine faculty. The advertisements described the study as “one of the first-ever fMRI studies that investigates how medical education shapes human brain function.” Non-physician participants were recruited via email advertisements distributed by department administrators and by word of mouth. The advertisements described the study as “an fMRI study on cognitive expertise” and emphasized that we were “especially interested in recruiting doctoral students/postdoctoral fellows” with expertise in their respective fields.

At the scanner, participants were told that the study “was about cognitive expertise.” No further information was provided until participants had completed the study and post-participation survey. In the post-participation study, we asked participants to answer several questions about their experience, including “what do you think the study was about?” Only one participant, a physician, identified that the study was about medical expertise (“Medical response to medical topics”). The rest of the participants described a range of general topics, including brain connectivity, problem solving, attention, decision making, and educational attainment.

## Appendix 4 Full description of experimental procedure

We used a “magic detection” task to encourage participants to process the meaning of the vignettes without making explicit causality judgments. Participants saw “magical” catch trials that closely resembled the experimental trials but were fantastical (e.g., “Sadie forgot to wash her face after she ran in the heat. Now she has a cucumber nose.”). On each trial, participants indicated via button press whether “something magical” occurred in the vignette (Yes/No). This semantic foil detection task encouraged participants to attend to the meaning of the critical vignettes while reading as naturally as possible. The use of an implicit task also helped prevent participants from discovering the experimental manipulation, as confirmed by a post-participation survey (see Appendix 3 for details).

Both sentences in each vignette were presented simultaneously for 7 s, one above the other, followed by a 12 s inter-trial interval. Each participant saw 38 trials per condition (152 trials) plus 36 “magical” catch trials (188 total trials) in one of two versions, counterbalanced across participants, such that individual participants did not see the same sentence in both causal and non-causal vignettes. The two stimulus versions had similar meanings but different surface forms (e.g., “Holly used a grimy faucet…” vs. “Steve used a dirty bathroom…”).

The experiment was divided into 6 10-minute runs containing 6-7 trials per condition per run presented in a pseudorandom order. Vignettes from the same experimental condition repeated no more than twice consecutively, vignettes that shared similar phrase structure never repeated within a run, vignettes that referred to the same illness never repeated consecutively, and vignettes from each condition, including catch trials, were equally distributed in time across the course of the experiment.

## Appendix 5 Full description of neural similarity analysis

In an exploratory analysis, we tested whether responses to illness causal inferences (*Illness-Causal*) in a temporoparietal network previously implicated in thinking about animates (i.e., PC, TPJ) were more similar within participant groups compared to across groups. Higher within-group neural similarity would suggest that medical training (or lack thereof) shapes neural responses to causal inferences about illness in similar ways. Following the results of univariate fROI analysis, we focused on the neural similarity of illness inferences in the left PC and left TPJ.

First, we first identified the top 300 vertices most responsive to illness causal inferences (*Illness-Causal > Mechanical-Causal*) in left PC and left TPJ group search spaces (Dufour et al., 2013) in individual participants. We then obtained one observation (z-scored beta parameter estimate of the individual-trial GLM) per trial of the *Illness-Causal* condition (n=38 trials). To reduce noise, we averaged data from pairs of trials with similar content, i.e., trials about the same illness (e.g., two trials about COVID-19), or if this was not possible, trials focusing on similar etiologies (e.g., two trials about developing different types of cancers due to genetic factors; see Supplementary Table 1 for a full list of pairings). One trial was excluded from this analysis (asthma) because no other trials were similar to it. To account for the resulting imbalance, in one instance three trials were averaged instead of two. Together, this resulted in 18 observations per participant, each representing the average of two (or three) similar *Illness-Causal* trials.

We then calculated the ranked similarity (Euclidean distance) between each pair of observations. To initially test whether there was any signal to be captured in our regions of interest across participants, we performed a leave-one-participant-out correlation procedure (Spearman’s rho), collapsing across groups. In this analysis, each individual participant’s ranked similarity values were compared to the average of the remaining participants’ average ranked similarity values. In the fROIs that exhibited average correlations greater than zero (one-tailed Student’s *t*-tests) we subsequently performed a more sensitive within- vs. across-group analysis. For each participant in a group, we correlated (Spearman’s rho) their ranked similarity values for illness causal inference trials with i) the average ranked similarities of the remaining participants in that group and ii) the average ranked similarity values of all participants in the other group. This yielded two correlation values (within-group and across-group) for each participant. We then performed a repeated-measures ANOVA to test for differences in within- vs. across-group similarity in both physicians and non-physicians.

## Appendix 6 Full behavioral results

For the mentalizing localizer experiment, there were no differences in accuracy or response time across groups in the comprehension question task (repeated measures ANOVA, condition (*mentalizing stories*, *physical stories*) x group (physicians, non-physicians), accuracy: main effect of group, *F*_(1,30)_ = 0.08, *p* = .78, group x condition interaction, *F*_(1,30)_ = 0.25, *p* = .62; response time: main effect of group, *F*_(1,30)_ = 0.26, *p* = .61, group x condition interaction, *F*_(1,30)_ = 0.26, *p* = .62). Across groups, there were no differences in accuracy across conditions (main effect of condition, *F*_(1,30)_ = 2.85, *p* = .1). Participants were faster to respond to comprehension questions about physical stories than mentalizing stories (main effect of condition, *F*_(1,30)_ = 18.46, *p* < .001).

For the language/logical reasoning localizer experiment, there were no group differences in accuracy (repeated measures ANOVA, condition (*language*, *logical reasoning*, *math*) x group (physicians, non-physicians), accuracy: main effect of group, *F*_(1,30)_ = 0.004, *p* = .95, condition x group interaction, *F*_(2,60)_ = 1.32, *p* = .28). There was a marginal effect in response time, whereby non-physicians responded marginally faster on the *logical reasoning* trials compared to physicians (response time: main effect of group, *F*_(1,30)_ = 0.51, *p* = .48, condition x group interaction, *F*_(2,60)_ = 3.06, *p* = .05). Across both groups, participants were significantly more accurate on *language* trials, followed by *math* and then *logical reasoning* (main effect of condition, *F*_(2,60)_ = 108.51, *p* < .001). Participant response time showed a similar pattern: *language* trials were completed the fastest, followed by *math* and then *logical reasoning* (main effect of condition, *F*_(2,60)_ = 95.74, *p* < .001).

## Appendix 7 Comparison of illness causal inferences and mechanical causal inferences in the FFA across groups (fROI analysis)

There was no difference between causal inferences about illness and mechanical breakdown in lateral ventral occipitotemporal cortex (i.e., fusiform face area) (individual-subject fROI leave-one-run-out analysis; repeated measures ANOVA, condition (*Illness-Causal*, *Mechanical-Causal*) x hemisphere (left, right) x group (physicians, non-physicians), main effect of condition, *F*_(1,30)_ = 0.02, *p* = .9). Responses to both conditions were stronger in the left hemisphere (main effect of hemisphere, *F*_(1,30)_ = 0.25, *p* = .62, condition x hemisphere interaction, *F*_(1,30)_ = 23.5, *p* < .001). No other significant effects were observed in this region (main effect of group, *F*_(1,30)_ = 3.17, *p* = .09, condition x group interaction, *F*_(1,30)_ = 0.5, *p* = .48, hemisphere x group interaction, *F*_(1,30)_ = 2.93, *p* = .1, condition x hemisphere x group interaction, *F*_(1,30)_ = 2.24, *p* = .15).

## Appendix 8 Comparison of illness causal inferences and mechanical causal inferences in the PC and TPJ across groups (fROI analysis)

Causal inferences about illness elicited left-lateralized neural responses in a temporoparietal semantic network previously implicated in knowledge about animate entities (e.g., people, animals). Precuneus (PC) and temporoparietal junction (TPJ) fROIs exhibited increased activity to causal inferences about illness compared to mechanical causal inferences (individual-subject fROI leave-one-run-out analysis; repeated measures ANOVA, condition (*Illness-Causal*, *Mechanical-Causal*) x hemisphere (left, right) x group (physicians, non-physicians), PC: main effect of condition, *F*_(1,30)_ = 27.45, *p* < .001, TPJ: main effect of condition, *F*_(1,30)_ = 11.41, *p* = .002). The difference between illness and mechanical causal vignettes was more pronounced in the left hemisphere (PC: main effect of hemisphere, *F*_(1,30)_ = 0.25, *p* = .62, condition x hemisphere interaction, *F*_(1,30)_ = 39.97, *p* < .001, TPJ: main effect of hemisphere, *F*_(1,30)_ = 4.78, *p* = .04, condition x hemisphere interaction, *F*_(1,30)_ = 9.10, *p* = .005).

In these same temporoparietal regions (i.e., the PC and TPJ), physicians exhibited higher responses to illness causal inferences compared to mechanical causal inferences (PC: main effect of group, *F*_(1,30)_ = 1.87, *p* = .18, condition x group interaction, *F*_(1,30)_ = 2.40, *p* = .13, hemisphere x group interaction, *F*_(1,30)_ = 0.13, *p* = .72, condition x hemisphere x group interaction, *F*_(1,30)_ = 6.42, *p* = .02; TPJ: main effect of group, *F*_(1,30)_ = 0.66, *p* = .42, condition x group interaction, *F*_(1,30)_ = 6.08, *p* = .02, hemisphere x group interaction, *F*_(1,30)_ = 2.19, *p* = .15, condition x hemisphere x group interaction, *F*_(1,30)_ = 5.92, *p* = .02). The difference between physicians and non-physicians was specific to the left hemisphere in both the PC and TPJ (repeated measures ANOVA, condition (*Illness-Causal*, *Mechanical-Causal*) x group (physicians, non-physicians), left PC: main effect of condition, *F*_(1,30)_ = 42.06, *p* < .001, main effect of group, *F*_(1,30)_ = 1.48, *p* = .23, condition x group interaction, *F*_(1,30)_ = 4.56, *p* = .04; right PC: main effect of condition, *F*_(1,30)_ = 8.74, *p* = .006, main effect of group, *F*_(1,30)_ = 1.97, *p* = .17, condition x group interaction, *F*_(1,30)_ = 0.36, *p* = .55; left TPJ: main effect of condition, *F*_(1,30)_ = 18.59, *p* < .001, main effect of group, *F*_(1,30)_ = 2.34, *p* = .14, condition x group interaction, *F*_(1,30)_ = 10.66, *p* = .003; right TPJ: main effect of condition, *F*_(1,30)_ = 1.78, *p* = .19, main effect of group, *F*_(1,30)_ = 0, *p* = .96, condition x group interaction, *F*_(1,30)_ = 0.72, *p* = .40; Figure 2).

## Appendix 9 Comparison of illness causal inferences and non-causal vignettes in the PC and TPJ across groups (fROI analysis)

Precuneus (PC) fROIs exhibited increased activity to causal inferences about illness compared to non-causal vignettes (individual-subject fROI leave-one-run-out analysis; repeated measures ANOVA, condition (*Illness-Causal*, combined *Noncausal-Illness First* + *Noncausal-Mechanical First*) x hemisphere (left, right) x group (physicians, non-physicians), main effect of condition, *F*_(1,30)_ = 13.78, *p* < .001). The difference between illness causal and non-causal vignettes was more pronounced in the left hemisphere (main effect of hemisphere, *F*_(1,30)_ = 2.36, *p* = .14, condition x hemisphere interaction, *F*_(1,30)_ = 20.02, *p* < .001). No group differences were observed (main effect of group, *F*_(1,30)_ = 2.60, *p* = .12, condition x group interaction, *F*_(1,30)_ = 1.07, *p* = .31, hemisphere x group interaction, *F*_(1,30)_ = 1.04, *p* = .32, condition x hemisphere x group interaction, *F*_(1,30)_ = 0.76, *p* = .39).

The TPJ showed a different pattern. Across the full sample, TPJ fROIs did not show increased activity to causal inferences about illness compared to non-causal vignettes (individual-subject fROI leave-one-run-out analysis; repeated measures ANOVA, condition (*Illness-Causal*, combined *Noncausal-Illness First* + *Noncausal-Mechanical First*) x hemisphere (left, right) x group (physicians, non-physicians), main effect of condition, *F*_(1,30)_ = 0.39, *p* = .54). However, we observed a significant three-way interaction between condition, hemisphere, and group, whereby only physicians exhibited increased activity to illness causal inferences compared to non-causal vignettes in the left TPJ (condition x hemisphere x group interaction, *F*_(1,30)_ = 5.51, *p* = .03, condition x group interaction, *F*_(1,30)_ = 5.72, *p* = .02, hemisphere x group interaction, *F*_(1,30)_ = 3.07, *p* = .09, condition x hemisphere interaction, *F*_(1,30)_ = 4.09, *p* = .05, main effect of group, *F*_(1,30)_ = 0.91, *p* = .35, main effect of hemisphere, *F*_(1,30)_ = 7.85, *p* = .009).

## Appendix 10 Spatial dissociation between responses to causal inferences about illness vs. mental states

Following our prior study (Hauptman & Bedny, 2025), we tested whether responses to illness inferences were located ventrally to mental state inference responses in a left PC search space (Dufour et al., 2013). In the current dataset, the z-coordinates of individual-subject activation peaks for illness inferences and mental state inferences were significantly different in both groups (repeated measures ANOVA, peak type (mental state inferences, illness causal inferences) x group (physicians, non-physicians), main effect of peak type, *F*_(1,30)_ = 34.85, *p* < .001, main effect of group, *F*_(1,30)_ = 1.43, *p* = .24, peak type x group interaction, *F*_(1,30)_ = 1.44, *p* = .24).

## Appendix 11 Comparison of mechanical causal inferences and illness causal inferences in the anterior PPA across groups (fROI analysis)

Anterior PPA fROIs exhibited increased activity to causal inferences about mechanical breakdown compared to illness causal inferences (individual-subject fROI leave-one-run-out analysis; repeated measures ANOVA, condition (*Illness-Causal*, *Mechanical-Causal*) x hemisphere (left, right) x group (physicians, non-physicians), main effect of condition, *F*_(1,30)_ = 31.12, *p* < .001). The difference between mechanical and illness causal vignettes was more pronounced in the left hemisphere (main effect of hemisphere, *F*_(1,30)_ = 1.58, *p* = .22, condition x hemisphere interaction, *F*_(1,30)_ = 8.61, *p* = .006). Effects of expertise were not observed in this region (main effect of group: *F*_(1,30)_ = 1.97, *p* = .17, condition x group interaction, *F*_(1,30)_ = 1.76, *p* = .19, hemisphere x group interaction, *F*_(1,30)_ = 0, *p* = .94, condition x hemisphere x group interaction, *F*_(1,30)_ = 0.02, *p* = .9).

## Appendix 12 Pre-registered analyses not included in the main text

### Individual-subject fROI analysis: univariate

Mentalizing fROIs were created by selecting the most mentalizing-responsive vertices (top 5%) in bilateral PC and TPJ search spaces (Dufour et al., 2013) using the *mentalizing stories* > *physical stories* contrast from the mentalizing localizer. We extracted PSC for each condition in the illness causal inference task compared to rest and compared the difference between *Illness-Causal* and *Mechanical-Causal* across participant groups (physicians, non-physicians) using repeated-measures ANOVAs.

The difference in activity between illness causal inferences and mechanical causal inferences was equivalent across groups in mentalizing PC and TPJ fROIs (repeated measures ANOVA, condition (*Illness-Causal*, *Mechanical-Causal*) x hemisphere (left, right) x group (physicians, non-physicians), PC: main effect of condition, *F*_(1,30)_ = 18.63, *p* < .001, main effect of group, *F*_(1,30)_ = 1.06, *p* = .31, condition x group interaction, *F*_(1,30)_ = 2.85, *p* = .1, condition x hemisphere interaction, *F*_(1,30)_ = 3, *p* = .09, hemisphere x group interaction, *F*_(1,30)_ = 0.18, *p* = .67, condition x hemisphere x group interaction, *F*_(1,30)_ = 2.60, *p* = .12; TPJ: main effect of condition, *F*_(1,30)_ = 6.25, *p* = .02, main effect of group: *F*_(1,30)_ = 0.05, *p* = .83, condition x group interaction, *F*_(1,30)_ = 3.40, *p* = .08, condition x hemisphere interaction, *F*_(1,30)_ = 0.13, *p* = .73, hemisphere x group interaction, *F*_(1,30)_ = 0.53, *p* = .47, condition x hemisphere x group interaction, *F*_(1,30)_ = 3.28, *p* = .08).

### Individual-subject fROI analysis: MVPA

We performed MVPA (PyMVPA toolbox; Hanke et al., 2009) in individual-subject fROIs to test whether patterns of activity in the PC distinguish between illness causal inferences and mechanical causal inferences. In each participant, we defined two fROIs in left PC search spaces (Dufour et al., 2013): 1) the top 300 vertices most responsive to mentalizing (*mentalizing stories* > *physical stories*, and 2) the top 300 vertices most responsive to causal inferences regardless of content (*Illness-Causal + Mechanical-Causal* > Rest).

For each vertex in each participant’s ROIs, we obtained one observation per condition (*Illness-Causal*, *Mechanical-Causal*) per run (z-scored beta parameter estimate of the GLM). A linear support vector machine (SVM) was then trained on data all but one of the runs and tested on the left-out run in a cross-validation procedure. Classification accuracy was averaged across all permutations of the training/test splits. We compared classifier performance within each fROI to chance (50%; one-tailed test). We also tested for an interaction between group and fROI. Significance was evaluated against an empirically generated null distribution using a combined permutation and bootstrap approach (Schreiber & Krekelberg, 2013; Stelzer et al., 2013). In this approach, t- and F-statistics obtained for the observed data are compared against an empirically generated null distribution. We report the t- and F-values obtained for the observed data and the nonparametric p-values, where p corresponds to the proportion of the shuffled analyses that generated a comparable or higher t/F value.

The null distribution was generated using a balanced block permutation test by shuffling condition labels within run 1000 times for each subject (Schreiber & Krekelberg, 2013). Then, a bootstrapping procedure was used to generate an empirical null distribution for each statistical test across participants by sampling one permuted accuracy value from each participant’s null distribution 15,000 times (with replacement) and running each statistical test on these permuted samples, thus generating a null distribution of 15,000 statistical values for each test (Stelzer et al. 2013).

A repeated-measures ANOVA revealed an effect of group, whereby physicians showed marginally higher decoding accuracies (*Illness-Causal* vs. *Mechanical-Causal*) across PC ROIs (repeated measures ANOVA, group (physicians, non-physicians) x fROI type (*mentalizing*, *causal inference*), main effect of group, *F*_(1,30)_ = 4.28, permuted *p* = .05). We also observed a significant group by fROI interaction, whereby higher decoding accuracies in mentalizing fROIs were observed in physicians (main effect of fROI type, *F*_(1,30)_ = 0.40, permuted *p* = .53, group x fROI type interaction, *F*_(1,30)_ = 5.71, permuted *p* = .02). A full list of results across fROIs and groups can be found in Supplementary Table 2.

### Searchlight MVPA

We used a linear support vector machine classifier (PyMVPA toolbox; Hanke et al., 2009) to test decoding between illness causal inferences and both non-causal conditions (*Illness-Causal* vs. *Noncausal-Illness First, Illness-Causal* vs. *Noncausal-Mechanical First*) across the whole cortex using a 10 mm radius spherical searchlight (according to geodesic distance, to better respect cortical anatomy over Euclidean distance; Glasser et al., 2013). This yielded for each participant two classification maps indicating the classifier’s accuracy in the neighborhood surrounding every vertex. Individual subject searchlight accuracy maps were averaged within group (physicians, non-physicians) for each analysis. Each of the non-physicians maps were then subtracted from the physicians maps to identify the regions where decoding was better in the physician group. We used a permutation approach to threshold this difference map by randomly shuffling the individuals assigned to each group and performing the same subtraction of averaged group maps (1000 times). This yielded an empirical null distribution of subtracted values for each vertex that was used to threshold the true group difference map at *p* < .05. We note that this correction procedure deviates from the procedure outlined in the pre-registration. We never carried out the pre-registered procedure: upon reaching the analysis stage, we felt that a permutation test of the group subtraction was more suitable than the pre-registered procedure, which involved correcting maps within groups separately and performing a single subtraction of the corrected maps.

Physicians showed higher decoding accuracy between *Illness-Causal* and *Noncausal-Illness First* in bilateral PC, left TPJ, left medial prefrontal cortex, bilateral insula, and right middle and inferior temporal gyrus. Physicians showed higher decoding accuracy between *Illness-Causal* and *Noncausal-Mechanical First* in left medial prefrontal cortex, left temporo-occipital cortex, and the right insula (Supplementary Figures 6, 12).

